# A glycan cluster on the SARS-CoV-2 spike ectodomain is recognized by Fab-dimerized glycan-reactive antibodies

**DOI:** 10.1101/2020.06.30.178897

**Authors:** Priyamvada Acharya, Wilton Williams, Rory Henderson, Katarzyna Janowska, Kartik Manne, Robert Parks, Margaret Deyton, Jordan Sprenz, Victoria Stalls, Megan Kopp, Katayoun Mansouri, Robert J Edwards, R. Ryan Meyerhoff, Thomas Oguin, Gregory Sempowski, Kevin Saunders, Barton F. Haynes

**Affiliations:** Duke Human Vaccine Institute, Durham NC 27710, USA; Duke University, Department of Surgery, Durham NC 27710, USA; Duke University, Department of Medicine, Durham NC 27710, USA; Duke University, Department of Immunology, Durham NC 27710, USA; Duke University, Division of Interventional Radiology, Department of Radiology, Durham NC 27710, USA

## Abstract

The COVID-19 pandemic caused by SARS-CoV-2 has escalated into a global crisis. The spike (S) protein that mediates cell entry and membrane fusion is the current focus of vaccine and therapeutic antibody development efforts. The S protein, like many other viral fusion proteins such as HIV-1 envelope (Env) and influenza hemagglutinin, is glycosylated with both complex and high mannose glycans. Here we demonstrate binding to the SARS-CoV-2 S protein by a category of Fab-dimerized glycan-reactive (FDG) HIV-1-induced broadly neutralizing antibodies (bnAbs). A 3.1 Å resolution cryo-EM structure of the S protein ectodomain bound to glycan-dependent HIV-1 bnAb 2G12 revealed a quaternary glycan epitope on the spike S2 domain involving multiple protomers. These data reveal a new epitope on the SARS-CoV-2 spike that can be targeted for vaccine design.

**Highlights:**

- Fab-dimerized, glycan-reactive (FDG) HIV-1 bnAbs cross-react with SARS-CoV-2 spike.
- 3.1 Å resolution cryo-EM structure reveals quaternary S2 epitope for HIV-1 bnAb 2G12.
- 2G12 targets glycans, at positions 709, 717 and 801, in the SARS-CoV-2 spike.
- Our studies suggest a common epitope for FDG antibodies centered around glycan 709.

## Introduction

The severe acute respiratory syndrome coronavirus-2 (SARS-CoV-2) emerged in late 2019 and rapidly spread, causing a global pandemic of a severe acute respiratory syndrome (COVID-19). The SARS coronavirus-2 (SARS-CoV-2) utilizes a heavily glycosylated spike (S) protein to bind its host receptor, angiotensin-converting enzyme 2 (ACE2), and mediate cell entry and fusion (*1*). The S protein is a trimeric class I fusion protein composed of two functional subunits responsible for receptor binding (S1 subunit) and membrane fusion (S2 subunit) (*2, 3*). The surface of the virally encoded S protein is covered by host-derived glycans with each trimer encoding 66 N-linked glycosylation sites (*4*).

Viral fusion proteins such as the spike proteins of diverse Coronaviruses, the HIV-1 envelope (Env) and influenza hemagglutinin (HA), are typically covered with high mannose and complex glycans (*5, 6*). Viruses rely on host glycosylation machinery to synthesize and express glycans, which cover the surface of proteins exposed on the virus and shield conserved neutralization epitopes from immune surveillance. While genetic and somatic diversities can preclude targeting protein epitopes for broad-spectrum neutralization, viruses may display common carbohydrate moieties that can be targeted by glycan binding agents. Indeed, targeting glycan moieties has been proposed as a general strategy for broad-spectrum virus neutralization (*7-9*). While one concern with this general concept is that the glycan targeting agents may have unfavorable cross-reactivity with human tissue by binding host carbohydrate moieties, heavily glycosylated viral proteins, of which HIV-1 Env is an example, have significant population of unprocessed oligomannose-type glycans that are not usually observed at high abundance on secreted mammalian glycoproteins (*10, 11*). This divergence from host cell glycosylation and the ‘non-self’ nature of these oligomannose glycans represent immunogenic targets, and indeed a number of anti-HIV-1 broadly neutralizing antibodies (bnAbs) have been isolated that specifically target gp120 glycans as part of their epitope (*12-14*).

Site-specific glycan analysis of the SARS-CoV-2 spike protein demonstrated 28% of oligomannose-type glycans on the S protein surface (*4*). Here, we raised the hypothesis that the glycan shield of the SARS-CoV-2 spike protein can be targeted by HIV-1 glycan-reactive antibodies. We tested SARS-CoV-2 S protein for binding by HIV-1 bnAb 2G12 that binds a glycan-only epitope on the HIV-1 Env (*12, 15, 16*), as well as by a panel of Fab-dimerized glycan-reactive (FDG) antibodies. We have recently demonstrated that FDG antibodies are common in the HIV-1 uninfected B cell repertoire, and they bound high mannose glycans on HIV-1 Env, yeast and other environmental antigens (*17*).

## Results

### Binding of FDG antibodies to SARS-CoV-2 spike

In a companion manuscript (*17*), we described highly prevalent FDG antibodies that target the high mannose glycan shield of the HIV-1 Env. Here, we test members of this category of antibodies for binding to the SARS-CoV-2 spike (S) protein. A SARS-CoV-2 S protein ectodomain construct incorporating residues 1-1208 of the SARS-CoV-2 S protein and harboring two stabilizing proline mutations in the S2 domain, a foldon trimerization domain, as well as C-terminal Twin Streptactin and His tags, were expressed and purified as described previously from HEK-293F cells (**Figure S1**) (*2*). We tested three different groups of FDG antibodies – (1) FDG antibodies that were isolated from SHIV-infected macaques and shown to broadly neutralize HIV-1 isolates (DH851.1 – DH851.3, and DH1003.2); (2) FDG precursor antibodies that were isolated from a HIV naïve human (DH1005), and SHIV infected (DH898.1 and DH898.4) or HIV Env vaccinated (DH717.1 – DH717.4, and DH501) macaques; and (3) antibody 2G12 that forms a subcategory on its own as the only variable heavy (VH) domain-swapped FDG antibody known to date. Antibody 2G12 is a bnAb that neutralizes heterologous HIV-1 isolates and binds to a conformational glycan cluster on HIV-1 Env (*15, 18, 19*).

To test whether the FDG antibodies that bind the HIV-1 Env glycan shield can also recognize the SARS-CoV-2 spike we performed ELISA and SPR assays (**Figures 1 and S2-S8**). As a positive control in the ELISA assays we included antibody D001 (see methods) that binds the receptor binding domain (RBD) of SARS-CoV-1 and SARS-CoV-2, and as a negative control, antibody CH65 (*20*) that binds the influenza hemagglutinin. We observed binding of the S protein to the RBD-directed antibody D001, whereas no binding was observed for negative control antibody, CH65. All FDG antibodies tested bound the SARS-Cov-2 spike to varying degrees **(Figures 1, S2 and S3**). To determine if these antibodies were targeting glycan epitopes on the SARS-CoV-2 S protein, we tested binding of the antibodies to the S protein in the presence of a high-mannose glycan (D-mannose) (**Figures 1, S2 and S3**). As expected, no inhibition was observed for the D001 antibody that binds the S protein RBD epitope on the spike protein. Both the non-VH domain-swapped DH851 clone antibodies (DH851.1 – DH851.3) that have heterologous HIV-1 bnAb activity, and the VH domain-swapped antibody 2G12, bound the SARS-CoV-2 S protein in a glycan-dependent manner (**Figures 1, S2 and S3**). Other rhesus FDG bnAb (DH1003.2) and precursor (DH898.4, DH717.1 – DH717.3, and DH501) antibodies, as well as human FDG precursor (DH1005), bound the spike protein and were inhibited by D-mannose to varying degrees. These results demonstrated specific binding of FDG antibodies for glycans on the SARS-CoV-2 S protein.

**Figure 1.**
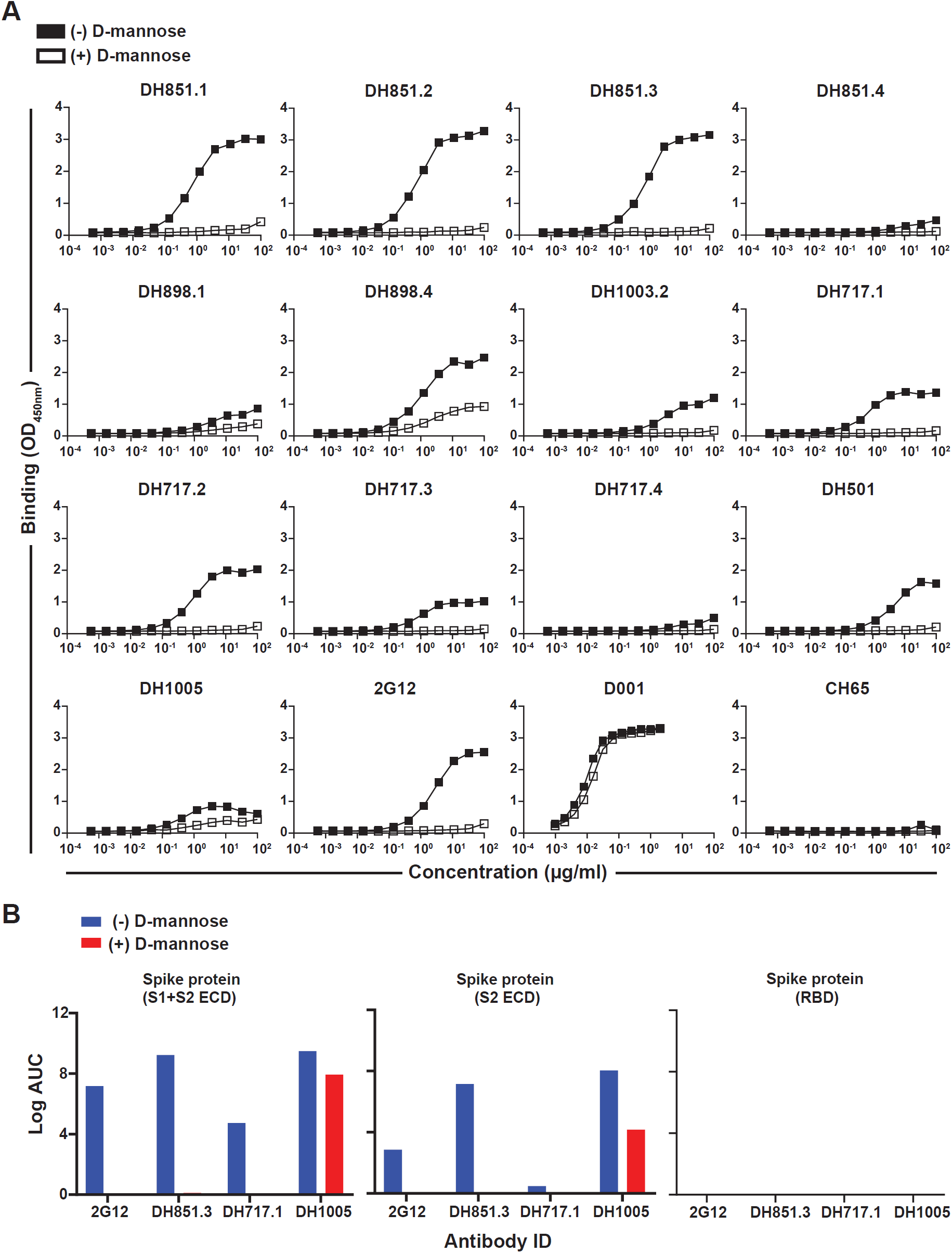
Glycan-dependent binding of FDG antibodies to SARS-CoV-2 spike protein. **(A)** We tested mAbs for binding to recombinant SARS-CoV-2 spike protein in ELISA. Antibody binding was assessed in the absence (-) or presence (+) of D-mannose [1M]. Serially-diluted mAbs were tested for binding (also see Figure S2 and S3). Mouse anti-rhesus or goat anti-human IgG-HRP secondary antibodies were used to detect binding by rhesus or human mAbs, respectively. Binding was measured at absorbance of OD_450nm_, whereas binding titers as Log area under the curve were reported in Figure S1. Control antibodies were SARS-CoV-1 RBD (D001) and influenza HA (CH65) mAbs. Data shown are from a representative assay. **(B)** Binding of FDG antibodies to a set of commercially available constructs expressing the SARS-CoV-2 S1 and S2 extracellular domain (left), S2 domain (middle), and the receptor binding domain (right). Blue and red bars show binding in the absence and presence of D-mannose [1M], respectively. Binding antibody titers are reported as area under the curve (AUC) and were calculated using Softmax software. Data shown are from a representative assay. All ELISAs (A-B) were done using BSA-based buffers (see methods).

To gain insight into the epitope on the SARS-CoV-2 spike ectodomain targeted by the FDG antibodies we chose 2G12 for structural determination in complex with the SARS-2 spike ectodomain. While the non-domain-swapped FDG antibodies exist as mixtures of Y- and I-shaped forms (*17*), the VH domain-swapped 2G12 presents a 100% I-shaped population that can be separated from its intermolecular dimer form by size exclusion chromatography (*21*) and digested using papain to yield a stable domain-swapped Fab dimer amenable to structural studies (**Figure S7**) (*12, 15, 18, 19*). The 2G12 Fab obtained by papain digestion of the 2G12 IgG (**Figure S7**) bound to SARS-CoV-2 spike with affinity of ∼313 nM (**Figure S8**), and on- and off-rates of 3.5 × 10^4^ M^-1^s^-1^ and 1.19 × 10^2^ s^-1^, respectively.

The FDG antibodies in Figure 1 as well as the HIV-1 bnAb 2G12 were tested in a SARS-CoV-2 plaque reduction neutralization assay and were all negative starting at 50 µg/ml (data not shown). Importantly, in none of the assays were enhanced numbers of viral plaques seen with the addition of FDG antibodies.

### Structure of SARS-CoV-2 spike bound to HIV-1 bnAb 2G12

To visualize the epitope of 2G12 bound to the SARS-CoV-2 spike we determined a cryo-EM structure of the 2G12 Fab bound to a stabilized SARS-CoV-2 ectodomain construct (*2*) (**Figures 2-6, S9 and S10; Table S1**). The cryo-EM dataset obtained from 6804 micrographs showed considerable compositional heterogeneity, revealing populations of unliganded spike, as well as spike bound to 2G12 at 1, 2 or 3 of it binding sites (**Figure S9**). The 2G12-bound spike was ∼15% of the total population. We obtained cryo-EM reconstructions of the SARS-CoV-2 S protein bound to a single 2G12 Fab at 3.3 Å from ∼ 108,000 particles, and S protein bound to two 2G12 Fabs at 3.2 Å resolution from ∼110,000 particles. (**Figure S9**). We also observed a smaller population (∼42,000 particles) that showed occupancy at all three binding sites, although the 2G12 density at one of the sites was substantially weaker than at the other two. Even upon extensive 3D classification of this population, the weaker density at the third site remained suggesting that this weak density was due to flexibility of the bound 2G12 rather than partial occupancy. To maximize the resolution of the 2G12 interface with the SARS-CoV-2 spike, we performed 3D refinement by combining all 2G12-bound particles and using a mask that focused the refinement on the S protein and one 2G12 Fab. On doing so, we obtained a 3.1 Å reconstruction of 2G12 Fab bound to the SARS-CoV-2 spike (**Figures 2 and S9; Table S1**). Fitting this map with the coordinates for the SARS-CoV-2 spike (PDB 6VXX) (*3*) and 2G12 Fab2 (PDB 6N35) (*12*), followed by iterative coordinate refinement and model building (*22, 23*) yielded a model for 2G12 Fab2 bound to the SARS-CoV-2 spike that was used for further analysis.

**Figure 2.**
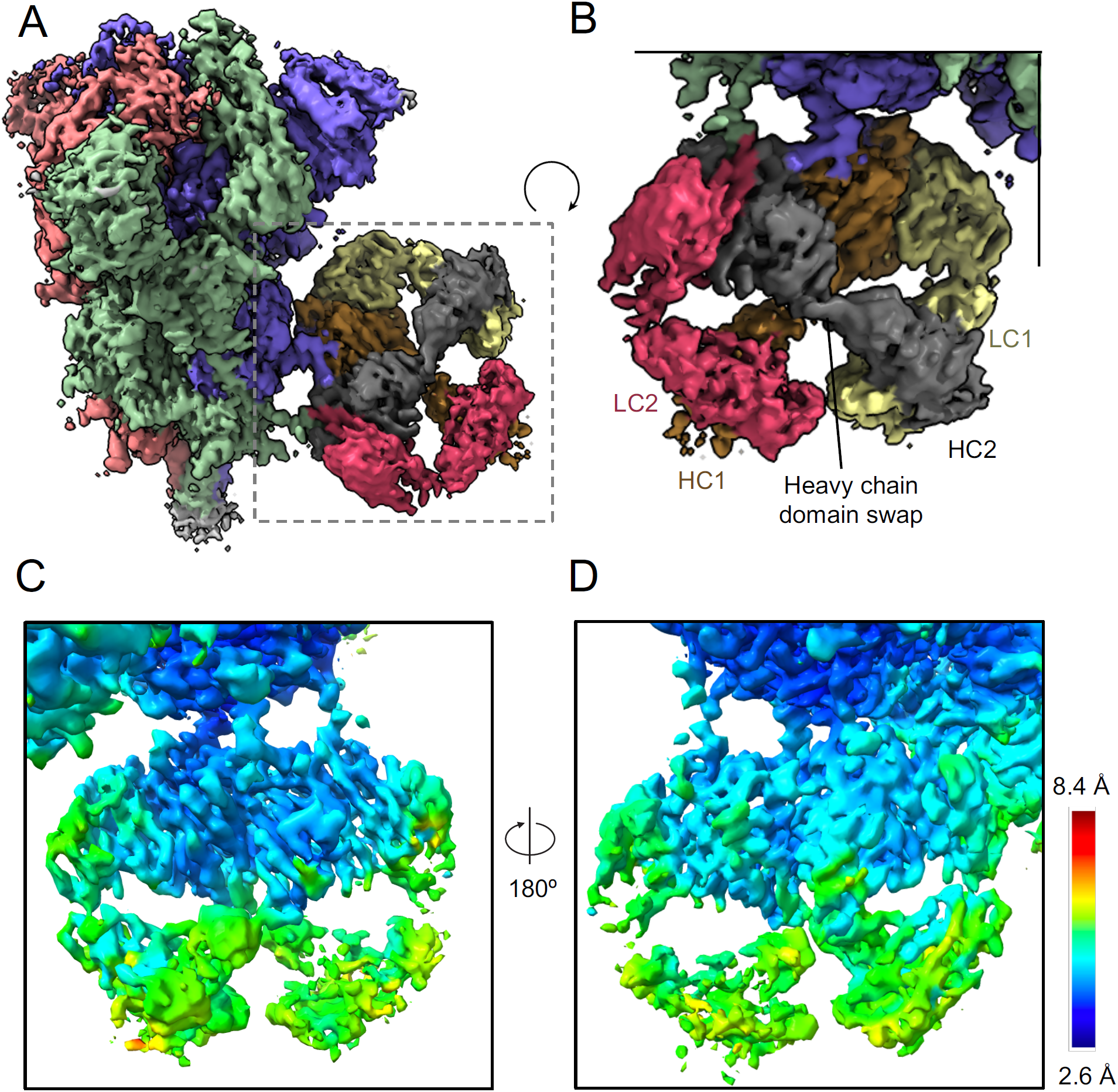
Cryo-EM structure of 2G12 Fab bound to the SARS-CoV-2 S protein. **(A)** Cryo-EM reconstruction of the SARS-CoV-2 spike bound to 2G12. The cryo-EM map is colored by chain. The SARS-CoV-2 S chains are colored salmon, green and blue, and the 2G12 chains are colored dark grey and orange for the heavy chains (HC), and yellow and dark pink for the light chains (LC). **(B)** Zoomed-in view of the bound 2G12 antibody, region marked with a dotted square in **(A). (C)** Zoomed-in view showing the cryo-EM reconstruction of the bound 2G12 Fab, colored by local resolution. **(D)** 180° rotated view of **(C)**.

The local resolutions of the cryo-EM reconstruction spanned a range from 2.6 Å – 8.4 Å (**Figures 2C-D, and S10**), with the lowest resolutions observed at the receptor binding domain, the N-terminal domain and the C-terminal ends of the SARS-CoV-2 S protein, and the constant domain of the 2G12 Fab. The binding interface of 2G12 with the SARS-CoV-2 was well resolved with the connecting glycans clearly visible in the electron density (**Figure 2**).

The VH domain swapped configuration of the 2G12 Fab-dimer (Fab2) was well-defined in the electron density of the bound 2G12 (**Figure 2A and B**). This unique domain-swapped architecture that has been described previously in a number of high-resolution structures solved using x-ray crystallography or cryo-EM (*12, 19, 24-27*), results in the creation of two secondary binding sites on either side of the VH/VH interface, in addition to the primary binding site at the VH/VL interface (*19*). The cryo-EM reconstruction of the Fab2 fragment of 2G12 bound to the SARS-Cov-2 spike revealed a quaternary epitope for 2G12 in the S2 subunit of the SARS-CoV-2 spike with glycans from different protomers clustered together to form the 2G12 epitope (**Figures 2-4**). Three S protein glycans were the primary contributors to the 2G12 binding epitope – glycans 717 and 801 from one protomer and glycan 709 from the adjacent protomer (**Figures 3-5**). The 2G12 Fab2 interaction with the SARS-CoV-2 spike buried a 1,500 Å^2^ area at its interface with the epitope composed entirely of N-linked glycans with the primary binding sites; each at one of the VH/VL interfaces, contacting glycans at positions N709 and N717 (**Figures 1A–1C**; see Figure S2 for glycan nomenclature). One of the two secondary binding sites created as a result of the VH/VH domain swapping contacted the glycan at positions N801 (Figures 3 and 4).

**Figure 3.**
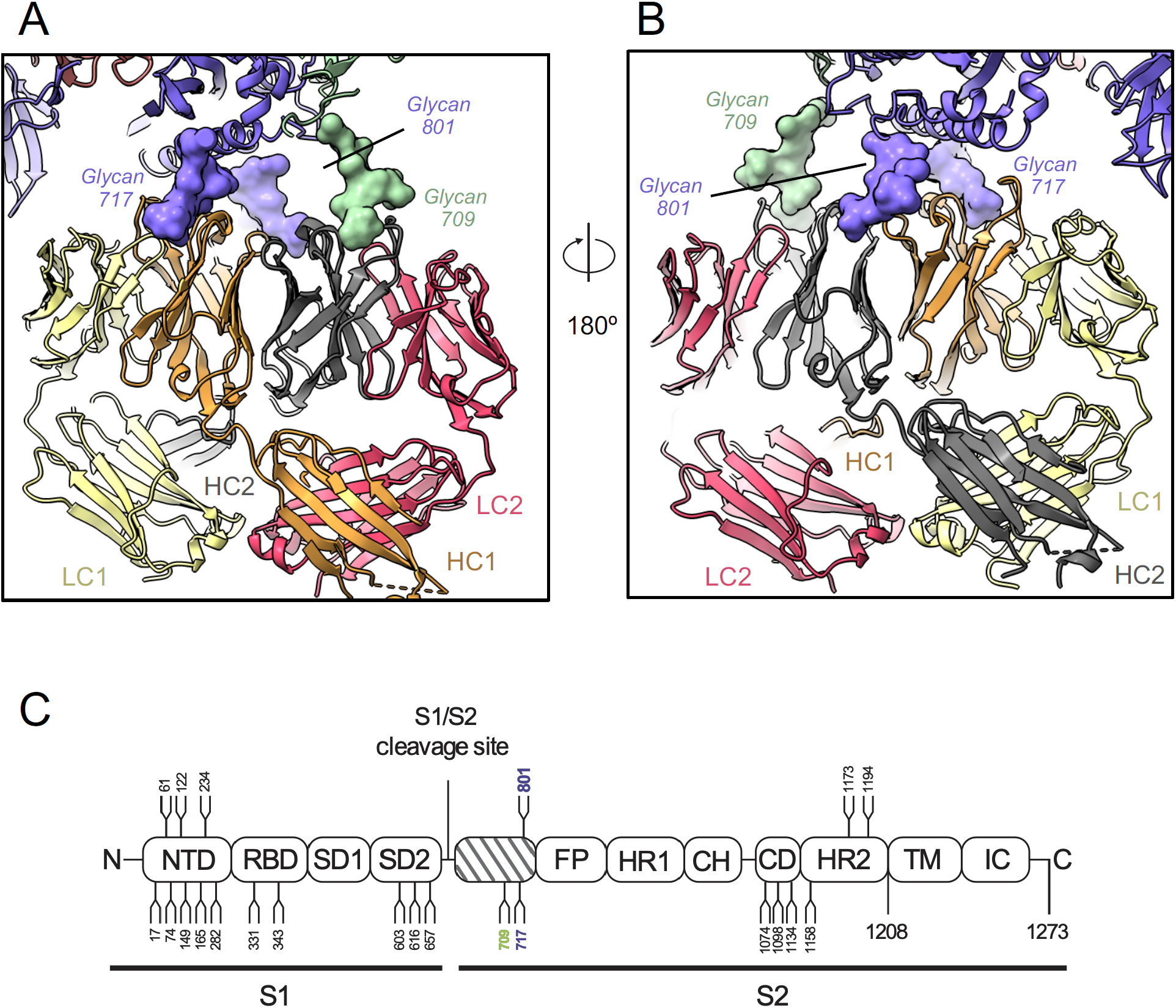
HIV-1 bnAb 2G12 binds a glycan-dominated, quaternary epitope on the SARS-CoV-2 S protein. **(A)** Zoomed-in view of domain-swapped, dimerized 2G12 Fab interacting with the SARS-CoV-2 S protein. The 2G12 epitope includes glycan 709 from one protomer and glycans 717 and 801 from an adjacent protomer. The structure is colored by chain following the same coloring scheme as in Figure 1A. 2G12 and the SARS-CoV-2 S are shown in cartoon representation. The interacting glycans are shown in surface representation. **(B)** 180° rotated view of **(A). (C)** Schematic showing the domain organization of the SARS-CoV-2 S protein. Positions of N-linked glycosylation sequons are numbered and shown as branches. The glycans making major contacts with 2G12 are colored by the protomer chain they belong to.

**Figure 4.**
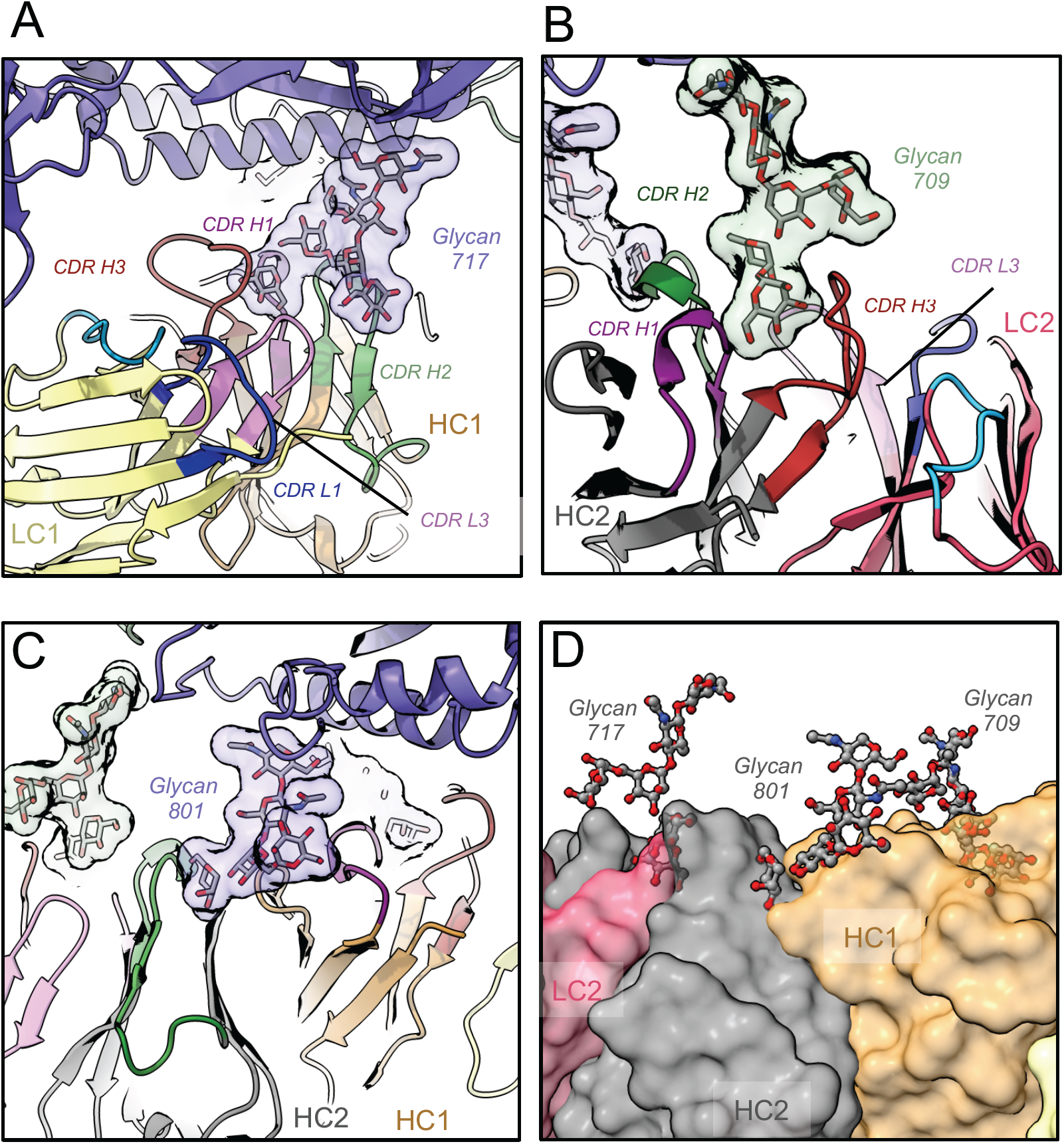
Details of 2G12 interactions with the SARS-CoV-2 S protein spike. **(A)** Binding site for SARS-CoV-2 glycan 717 at the pocket formed by the CDR H1, CDR H2, CDR H3 and CDR L3 loops in one of the Fab moieties **(B)** Binding site for SARS-CoV-2 glycan 709 at the pocket formed by the CDR H1, CDR H2, CDR H3 and CDR L3 loops in the other Fab moiety. **(C)** Binding site for SARS-CoV-2 glycan 801 at the 2G12 VH/VH domain swap interface **(D)** Surface representation of the 2G12 paratope, with 2G12 shown in surface representations and the interacting glycans in ball and stick representation.

### Identification of a common epitope on the SARS-CoV-2 S protein recognized by Fab-dimerized glycan-reactive antibodies

Because glycan-only epitopes are expected to be more mobile than a protein epitope we investigated the extent of disorder and motion in the position of the 2G12 bound to the S protein. Heterogeneous 3D classification performed on the cryo-EM dataset using a mask including the 2G12 density and S2 density, and excluding the S1 and the HR2 regions, revealed two distinct 2G12 orientations that were related by a 9-degree angular displacement about a hinge at glycan 709 (**Figure 5A and B**). To test the importance of this residue that appeared to be a pivot point for the bound 2G12, a N709A glycan deletion mutant was expressed and purified, and confirmed by NSEM to contain well-dispersed and properly folded spike particles (**Figure S1**). Deletion of the glycan at position 709 resulted in abrogation of 2G12 Fab binding, thus confirming that glycan 709 is critical for the binding of 2G12 to the SARS-Cov-2 spike (**Figure 5C**).

**Figure 5.**
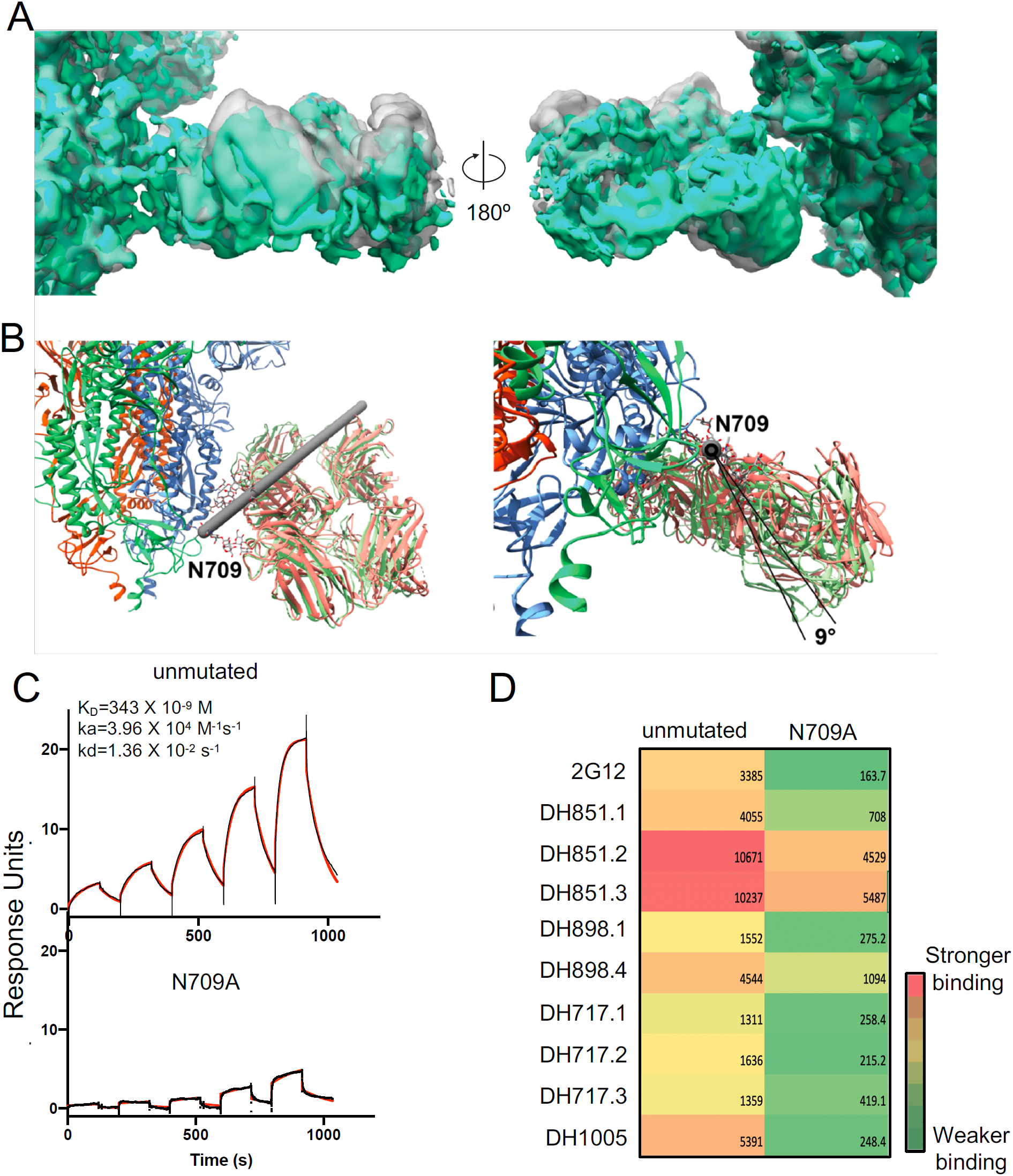
Heterogeneity and motion of the bound 2G12. **(A)** Two distinct states were resolved from the cryo-EM data by heterogeneous classification. Density for the two observed states are shown in green and grey. **(B)** Cartoon representation of the SARS-CoV-2 S-protein (bright green, bright orange, blue) and the two 2G12 orientations. The axis of rotation is represented by a grey cylinder. **(C)** Binding of 2G12 Fab2 to (top) unmutated spike and (bottom) N709A mutant. Binding of 2G12 Fab2 to the SARS-CoV-2 spike constructs was measured by SPR using single-cycle kinetics. The black lines show the data and the red lines the fit to a 1:1 Langmuir binding model. **(D)** Binding of the unmutated spike and the N709A mutant to a panel of FDG antibodies measured by SPR and shown as a heatmap. The spikes were captured via their C-terminal Twin Streptactin tags at ∼1000 RU levels on two alternate flow cells of a streptavidin coated chip. 200 nM of each FDG antibody was flowed over all four flow cells and the binding curves were double-reference subtracted (see also Figure S11). Areas under the curve were calculated in GraphPad Prism and are listed.

We next tested the binding of the panel of non-domain-swapped FDG antibodies to the N709A glycan-deleted mutant in an SPR assay. The N709 glycan deletion either abrogated or substantially reduced the binding of the FDG IgGs tested (**Figures 5D and S11**), thus suggesting a common epitope on the SARS-CoV-2 spike that is recognized by FDG antibodies.

### Comparison of 2G12 binding to SARS-CoV-2 spike versus HIV-1 Env

The VH domain-swapped 2G12 has been shown to bind isolated Man9 glycans, and to HIV-1 Env. In this study we demonstrate that 2G12 can also bind the SARS-CoV-2 S protein (**Figure 6A-D**) (*12, 19*). While 2G12 can recognize free mannose in its non-domain-exchanged form, it only binds the HIV-1 glycan shield if domain-exchanged, thus highlighting the importance of avidity for its interactions with glycan clusters (*27*). The structure of 2G12 bound to HIV-1 Env revealed binding sites for 4 HIV-1 Env glycans on 2G12 (*15, 18*), one each at the two primary binding sites at the VH/VL interface and the other two at the secondary sites at the VH/VH interface created as a result of domain-swapping (**Figures 6B**). Consistent with its glycan-only epitope, 2G12 binding to a soluble HIV-1 Env construct is inhibited by D-mannose (**Figure 6D**). In the structure of 2G12 Fab2 bound to the SARS-CoV-2 S protein, only 3 of these glycan binding sites on 2G12 were occupied by spike glycans (**Figure 6C**) and, while 2G12 showed robust glycan-dependent binding to the SARS-CoV-2 S protein, the binding was weaker than for the HIV-1 Env (**Figure 6D**). Another noteworthy difference between the binding of 2G12 to HIV-1 Env versus is binding to SARS-CoV-2 S is that on the HIV-1 Env all the binding glycans come from a single gp120 monomer in the HIV-1 Env trimer, whereas in the SARS-CoV-2 S protein the binding site is quaternary and the 2G12-binding glycans at each site are contributed by two of the three protomers, making this one of the first examples of a quaternary epitope spanning multiple protomers on the SARS-CoV-2 S protein (*28*), and the first in the S2 domain.

**Figure 6.**
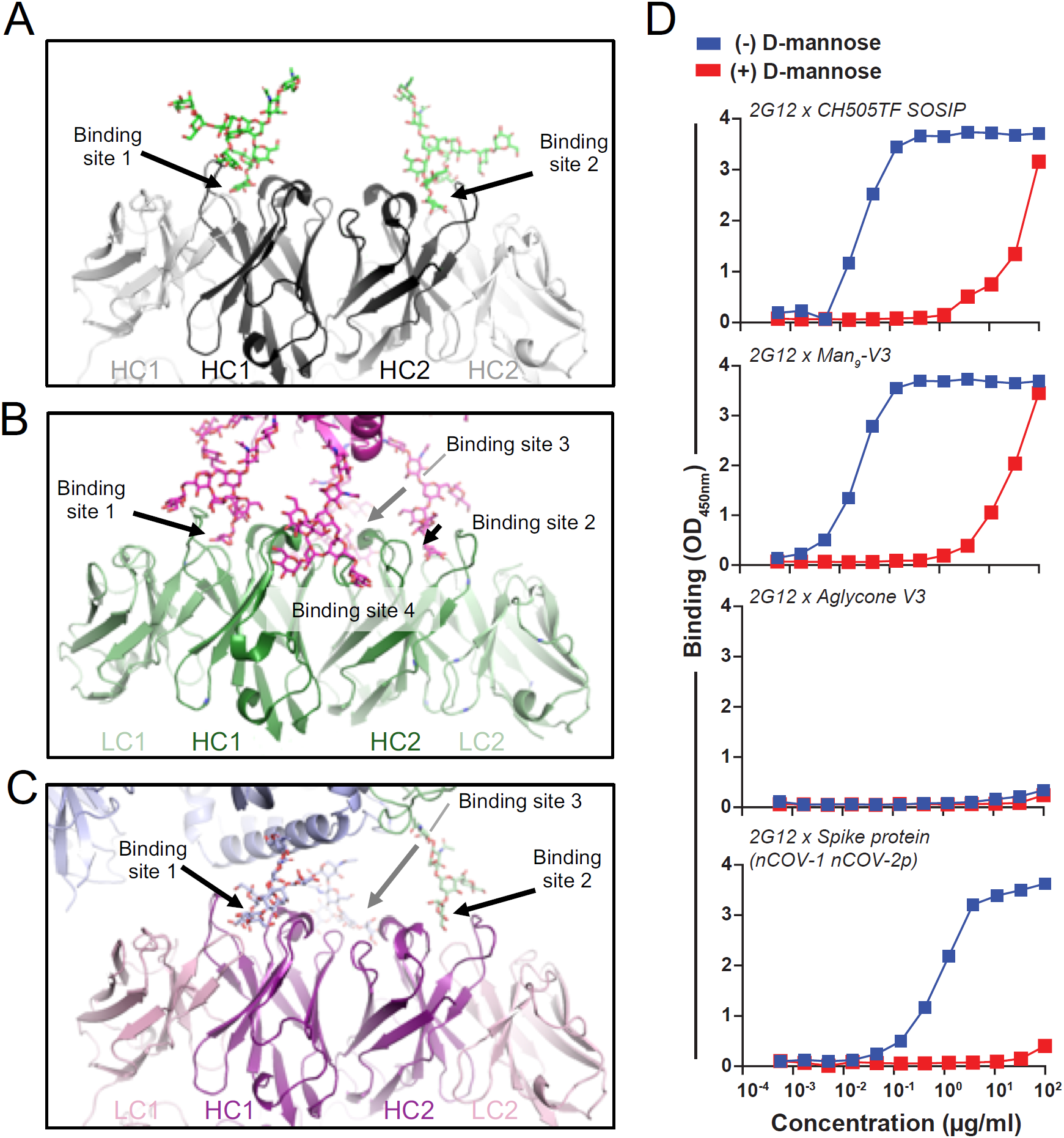
FDG antibody 2G12 leverages avidity to interact with glycan clusters on HIV-1 Env and SARS-CoV-2 S spike. **(A)** Binding of 2G12 to Man9 (PDB: 6N2X). Heavy chains are colored black, light chains are colored white, glycans are shown as sticks and colored by atom with carbons colored green. **(B)** Binding of 2G12 to HIV-1 Env (PDB: 6OZC). Heavy chains are colored dark green, light chains are colored light green, glycans are shown as sticks and colored by atom with carbons colored megante. **(C)** Binding of 2G12 to SARS-CoV-2 spike (this study). Heavy chains are colored deep purple, light chains are colored pink, glycans are shown as sticks and colored by atom with carbons colored slate blue for glycans coming from one protomer and pale green for glycans originating from the adjacent protomer **(D)** Binding of 2G12 to (from top to bottom) HIV-1 CH505TF SOSIP, Man_9_-V3 glycopeptide, Aglycone V3, and SARS-CoV-2 spike ectodomain. HIV-1 CH505TF SOSIP was captured using mouse anti-AVI-tag mAb, whereas SARS-CoV-2 ectodomain and peptides (Man_9_-V3 and Aglycone) were captured using streptavidin. Blue and red symbols indicate binding in the absence and presence of D-mannose [1M], respectively. Binding was measured at absorbance of OD_450nm_. Data shown are from a representative assay using BSA-based buffers (see methods).

## Discussion

In this paper we demonstrated that a category of prevalent glycan-reactive antibodies, termed FDG antibodies, can bind to high mannose glycans on the surface of the SARS-CoV-2 spike protein. FDG antibodies can bind to isolated high mannose glycans (*17, 29*) as well as high mannose glycans on HIV-1 and on fungal pathogens such as *Candida albicans* and *Cryptococcus neoformans* (*17, 30*). With HIV-1 infection in humans or SHIV infection of macaques, FDG antibodies can be affinity matured to achieve heterologous breadth. While the 2G12 HIV-1 broadly neutralizing antibody has VH domain swapped architecture (*12*), this architecture was so unique that 2G12 was the only antibody of its kind known for a long time. However, we recently discovered non-domain-swapped FDG antibodies that dimerize without VH domain swapping and are capable of broad HIV-1 broad neutralization, as shown for the DH851 IgG clonal lineage antibodies (*17*). Here we show, in addition to 2G12 binding, that VH domain swapping is also not required for FDG antibody recognition of the SARS-CoV-2 spike protein as demonstrated by its binding to the non-domain-swapped FDG antibodies. While the HIV-1 Env-reactive FDG antibodies tested in this study do not neutralize SARS-CoV-2, the demonstration that a common epitope for FDG antibodies exists on the surface of the SARS-CoV-2 spike, raises the hypothesis that SARS-CoV-2 infection may enjoin FDG precursors and select for FDG affinity matured antibodies that evolve the capacity to neutralize SARS-CoV-2 via glycan recognition.

That FDG precursors are common in the human B cell repertoire demonstrates that they are not deleted due to immune tolerance mechanisms (*17*). It has been previously postulated that the 2G12 and other glycan-reactive HIV-1 antibodies are rare due to reactivity with self-glycans (*31*). In this regard, select HIV-1 glycan reactive antibodies have been shown to react with the surface of immune cell subsets (*32*). Whether immune tolerance mechanisms will limit the affinity maturation of FDG antibodies to potently neutralize SARS-CoV-2 is yet unknown.

These data along with the companion paper describing the category of FDG anti-HIV-1 antibodies raise the hypothesis that preexisting antibodies raised by high mannose-containing environmental antigens such as fungi or host molecules can be co-opted by HIV-1 Env to become HIV-1 broadly neutralizing antibodies. Here we propose this same set of FDG precursor antibodies might be able to be targeted by the S protein of SARS-CoV-2 in the setting of COVID-19 disease or vaccination and induce SARS-CoV-2 neutralizing antibodies. It will be of interest to determine if FDG category of S-protein-reactive antibodies that can neutralize SARS-CoV-2 can be isolated from COVID-19 infected or vaccinated individuals. In this regard, Ng *et al* have demonstrated pre-existing S2 IgG antibodies to SARS-CoV-2 S-protein likely induced by human CoVs that cause common cold syndromes. Some of these sera were able to neutralize SARS-CoV-2 (*33*). However, it remains unknown if such pre-existing S2-protein antibodies are glycan reactive.

In summary, we demonstrate the reactivity of a pool of glycan-reactive antibodies with a quaternary epitope on the SARS-CoV-2 spike protein. A key study will be to determine if FDG antibodies can be recruited by vaccination to develop into antibodies that can play a preventive role in protection from SARS-CoV-2 infection.

## Acknowledgements

Initial cryo-EM data were collected on the Titan Krios at the Shared Materials and Instrumentation Facility in Duke University. High resolution cryo-EM data were collected at the National Center for Cryo-EM Access and Training (NCCAT) and the Simons Electron Microscopy Center located at the New York Structural Biology Center, supported by the NIH Common Fund Transformative High Resolution Cryo-Electron Microscopy program (U24 GM129539,) and by grants from the Simons Foundation (SF349247) and NY State. We thank Ed Eng, Daija Bobe, Mark Walters and Holly Leddy for microscope alignments and assistance with cryo-EM data collection. This study utilized the computational resources offered by Duke Research Computing (http://rc.duke.edu) at Duke University. We thank M. DeLong, C. Kneifel, M. Newton, V. Orlikowski, T. Milledge, and D. Lane from the Duke Office of Information Technology and Research Computing for helping set up and maintain the computing environment. The following reagent was deposited by the Centers for Disease Control and Prevention and obtained through BEI Resources, NIAID, NIH: SARS-Related Coronavirus 2, Isolate USA-WA1/2020, NR-52281.SARS-CoV-2 neutralization assays were performed under BSL3 in the Duke Regional Biocontainment Laboratory which received partial support for construction from NIH/NIAID (UC6AI058607). This work was supported by Translating Duke Health Initiative, NIH NIAID extramural project grant R01 AI145687 and a contract from the State of North Carolina Pandemic Recovery Office through funds from the Coronavirus Aid, Relief, and Economic Security (CARES) Act.

## Author contributions

P.A. and B.F.H. conceived and designed the study, evaluated all data, supervised the study, and wrote the paper. P. A. designed and performed the structural studies, and SPR assays. W.B.W. isolated and characterized FDG antibodies from macaque SHIV infections and HIV Env vaccinations, and HIV naïve humans, designed and performed ELISA assays, analyzed data, co-wrote, and edited the paper. R.C.H. performed structural analysis. K.J. performed SPR analysis and purified proteins. K.Manne purified proteins, and performed coordinate refinement and structural analysis. V.S. purified proteins. M.D., J.S. and R.P. characterized antibody binding specificities. K. Mansouri and R.J.E. performed NSEM experiments. R.R.M. and K.O.S. provided key reagents. T.O. and G.D. performed neutralization assays. M.K. performed specimen optimization and grid screening.

## MATERIALS AND METHODS

### Expression of recombinant SARS-CoV-2 spike

The SARS-CoV-2 ectodomain constructs were produced and purified as described previously (*2*). Briefly, a gene encoding residues 1-1208 of the SARS-CoV-2 S (GenBank: MN908947) with proline substitutions at residues 986 and 987, a “GSAS” substitution at the furin cleavage site (residues 682–685), a C-terminal T4 fibritin trimerization motif, an HRV3C protease cleavage site, a TwinStrepTag and an 8XHisTag was synthesized and cloned into the mammalian expression vector pαH. All mutants were introduced in this background. expression plasmids encoding the ectodomain sequence were used to transiently transfect FreeStyle293F cells using Turbo293 (SpeedBiosystems). Protein was purified on the sixth day post transfection from the filtered supernatant using StrepTactin resin (IBA) (Figure S1). Affinity purified protein was run over a Superose 6/300 increase column. The purified protein was flash-frozen in liquid nitrogen and stored at -80° C.

### Expression of antibodies

Rhesus FDG mAbs DH851.1 – DH851.3, DH1003.2, DH898.1 and DH898.4, DH717.1 – DH717.4, and DH501 and human FDG antibody DH1005 were expressed as previously described (*17*). Antibody 2G12 was obtained from three sources, either produced from two sources as described from the original sequence (*17*) 2G12_G1M17 or 2G12-rV2, or purchased from Polymun, Vienna Austria.

### Glycan-dependent binding of FDG mAbs to SARS-CoV-2 spike protein

Recombinant Fab-dimerized glycan (FDG)-reactive monoclonal antibodies (mAbs) were tested for binding SARS-CoV-2 trimer (lot 62KJ) in ELISA in the absence or presence of single monomer D-mannose sugar. SARS-CoV-2 trimeric spike protein was captured by streptavidin to individual wells of a 384-well plate, and serially diluted mAbs were tested for binding. ELISA binding assays were optimized to use SARS-CoV-2 spike protein that was aliquotted and frozen upon production; desired aliquots of frozen spike protein were thawed once at 37° and stored overnight at room temperature. SARS-CoV-2 spike protein, and Man_9_-V3 and Aglycone V3 peptides were captured via streptavidin on Nunc-absorb ELISA plates using PBS-based buffers and assay conditions as previously described (*34, 35*). HIV-1 CH505TF SOSIP trimer and commercially-obtained constructs of SARS-CoV-2 spike ectodomain (S1+S2 ECD, S2 ECD and RBD) (Sino Biological Inc cat# 40589-V08B1 and 40590-V08B respectively and RBD from Genescript cat# Z03483) were captured using mouse anti-AVI-tag mAb (Avidity LLC, Aurora, CO). In brief, we coated 30ng of streptavidin in 15µl at 2µg/ml or 30 ng of mouse anti-AVI-tag mAb in 15µl at 2µg/ml per well of a 384-well Nunc-absorb ELISA plate, sealed and incubated overnight at 4°C. SARS-CoV-2 spike protein (2µg/ml) were added to streptavidin in 10µl per well of a 384-well plate for one hour at room temperature to facilitate protein capture. HIV-1 CH505TF SOSIP trimer and SARS-CoV-2 S1+S2 ECD, S2 ECD and RBD spike proteins (2 µg/ml) were added to anti-AVI mAb in 10µl per well of a 384-well plate for one hour at room temperature to facilitate protein capture. Mouse anti-monkey IgG-HRP (Southern Biotech, CAT# 4700-05) or Goat anti-human IgG-HRP (Jackson ImmunoResearch Laboratories, CAT# 109-035-098) secondary antibodies were used to detect mAb bound to the SARS-CoV-2 spike protein. HRP detection was subsequently quantified with 3,3′,5,5′-tetramethylbenzidine (TMB) by measuring binding levels at an absorbance of 450nm; binding titers were reported as Log area under the curve (AUC). Commercially obtained D-mannose (Sigma, St. Louis, MO) was used to outcompete mAb binding to glycans on SARS-CoV-2; D-mannose solutions were produced in ELISA buffers. Competition ELISAs with D-mannose were performed in two separate assays; each with [0.5M] (data not shown) or [1.0M] D-mannose. Mouse-Human chimeric mAb D001 (SARS-CoV RBD mAb; Sino Biological Inc Cat# 40150-D001) was used as a control mAb; D001 was tested at 2µg/ml and 2-fold serial dilutions (10x), in contrast to all other mAbs tested at 100µg/ml and 3-fold serial dilutions (10x). Anti-influenza CH65 mAb was used a negative control mAb. We found that goat serum containing buffer (Superblock) – PBS, 4% (w/v) whey protein (BiPro USA), 15% normal goat serum (Invitrogen), 0.5% Tween 20, and 0.05% sodium azide (Sigma-Aldrich) – inhibited binding of glycan-dependent FDG mAbs to SARS-CoV-2 spike protein, whereas positive control RBD-binding mAb D001 bound the spike protein in the presence of superblock. ELISAs using superblock were previously described (*36*). Moreover, whereas the 2G12 from Polymun bound to S protein in surface plasmon reasonance, it was less potent than the recombinantly expressed 2G12 antibodies, and did not bind well in ELISA. Commercial 2G12 from Polymun is provided in a maltose (mannose) buffer.

### Neutralization assays

The neutralization assays were performed using SARS-CoV-2 isolate USA-WA1/2020, RVU Lot: NR52281-20200314. Source material supplied by BEI resources, catalog number NR-52281. Data are reported as PRNT_50_ or 50% reduction in foci count relative to back titer input verification. Endpoint titer of a given dilution series is reported as the concentration of the lowest continuous dilution with a foci count less than or equal to the 50% neutralization cutoff. All experimental samples were assayed as at least duplicate dilution series. Virus input and internal assay controls were within accepted tolerances.

### Negative-stain electron microscopy

A 100 µg/ml final concentration of the spike was made in 1:1 ratio of 0.15% glutaraldehyde in HBS pH 7.4 (20mM HEPES, 150mM NaCl) to 10% Glycerol in HBS, pH 7.4. After 5 min incubation Tris pH 7.4 was added from a 1M stock to a final concentration of 0.075M to quench the glutaraldehyde and incubated for 5min. The carbon coated grids (CF300-cu, EMS) were glow discharged for 20sec at 15mA. A 5 µl of sample incubated on grid for 10-15 sec, blotted and then stained with 2% uranyl formate. The antibodies were diluted in 0.02% Ruthenium red in HBS pH 7.4. After 5-10min incubation, a 5 µl droplet loaded on a glow discharged carbon coated grid, briefly rinsed with dH_2_O and then stained with 2% uranyl formate. Images were obtained with a Philips EM420 electron microscope operated at 120 kV, at 82,000× magnification and a 4.02 Å pixel size. The RELION program was used to perform class averaging of the single-particle images.

### Surface Plasmon Resonance (SPR)

The binding FDG antibodies to SARS-CoV-2 spike was assessed by surface plasmon resonance on Biacore T-200 (GE-Healthcare) at 25°C with HBS-EP+ (10 mM HEPES, pH 7.4, 150 mM NaCl, 3 mM EDTA, and 0.05% surfactant P-20) as the running buffer. Binding was measured in two different formats. In the first format (Figures S4 and S5A), SARS-CoV-2 spike was captured on a SA chip and binding response was measured by flowing over IgG solutions. The surface was regenerated between injections by flowing over 1M NaCl in 50mM NaOH solution for 10s with flow rate of 100µl/min. In the second format (Figure S5B, S6, S11) IgG were captured on an anti-Fc sensor surface and binding was assessed by flowing over solutions of SARS-CoV-2 in running buffer. The surface was regenerated between injections by flowing over 3M MgCl_2_ for 10s with flow rate of 100µl/min.

For determining affinity and kinetics of 2G12 interaction with SARS-CoV-2 spike, single cycle kinetics analysis was performed on the SARS-CoV-2 S protein immobilized on a streptavidin (SA) chip with five injections of 2G12 Fab, prepared by digestion with papain as described previously (*18*), at 3.125 nM, 6.25 nM, 12.5 nM, 25 nM and 50 nM concentrations. The data were fit to a 1:1 Langmuir binding model.

### Cryo-electron microscopy

#### Cryo-EM sample preparation

Purified SARS-CoV-2 spike preparations were diluted to a final concentration of ∼1 mg/mL in 2 mM Tris pH 8.0, 200 mM NaCl and 0.02% NaN3, were mixed with 6-fold molar excess of 2G12 Fab and incubated for 2 hours at room tempearture. 2.5 uL of protein was deposited on a CF-1.2/1.3 grid that had been glow discharged for 30 seconds in a PELCO easiGlow™ Glow Discharge Cleaning System. After a 30 s incubation in >95% humidity, excess protein was blotted away for 2.5 seconds before being plunge frozen into liquid ethane using a Leica EM GP2 plunge freezer (Leica Microsystems).

#### Cryo-EM data collection

Cryo-EM imaging was performed on a FEI Titan Krios microscope (Thermo Fisher Scientific) operated at 300 kV. Data were acquired with a Gatan K3 detector operated in counting mode using the Leginon system. The K3 experimental parameters are 2500 ms exposure with 50 frames at 50 ms framerate with a total dose of 66.43 e-/A2 and a pixel size of ∼1.058Å/px. This system was also energy-filtered with a slit width of 20 eV. A total of 6804 images were collected.

#### Cryo-EM data processing

Cryo-EM image quality was monitored on-the-fly during data collection using automated processing routines. Individual frames were aligned and dose-weighted using MotionCor2 (*37*) implemented within the Appion pipeline (*38*). Data processing was performed within cryoSPARC (*39*) including particle picking, multiple rounds of 2D classification, *ab initio* reconstruction, heterogeneous and homogeneous map refinements, and non-uniform map refinements. Heterogenous classifications were performed within RELION (*40*) using masks produced in Chimera.

### Cryo-EM model fitting

Structure of the all ‘down’ state (PDB ID 6VXX) the previously published SARS-CoV-2 ectodomain, and a structure of the 2G12 Fab bound to Man1-2 (PDB ID 6N35) were used to fit the cryo-EM maps in ChimeraX. Coordinates were then fit manually in Coot (*23*) following iterative refinement using Isolde (*22*) and subsequent manual coordinate fitting in Coot. Structure and map analysis was performed using PyMol and ChimeraX (*41*).

### Data availability

The cryo-EM maps and fitted coordinates are in the process of being deposited to the electron microscopy database (EMDB) and the RCSB Protein Data Bank, respectively. The sequences of the FDG antibody heavy and light chain genes were deposited in GenBank as previously described (*17, 29*)

---

**Table S1:** Cryo-EM Data Collection and Refinement Statistics.

|  | 2 2G12-bound<br>SARS-CoV-2 S | 1 2G12-bound<br>SARS-CoV-2 S | 1 2G12-bound<br>SARS-CoV-2 S<br>(focused<br>refinement) |
| --- | --- | --- | --- |
| Data Collection |  |  |  |
| Microscope | FEI Titan Krios |  |  |
| Voltage (kV) | 300 |  |  |
| Electron dose (e<sup>-</sup>/Å<sup>2</sup>) | 66.43 |  |  |
| Detector | Gatan K3 |  |  |
| Pixel Size (Å) | 1.058 |  |  |
| Defocus Range (µm) | 0.57-2.92 |  |  |
| Magnification | 81000 |  |  |
| Micrographs Collected | 6804 |  |  |
| Reconstruction |  |  |  |
| Software | cryoSPARC | cryoSPARC | cryoSPARC |
| Particles | 109,774 | 107,551 | 222304 |
| Symmetry | C1 | C1 | C1 |
| Box size (pix) | 320 | 320 | 320 |
| Resolution (Å)<sup>\$</sup> | | | |
| Corrected | 3.3 | 3.2 | 3.1 |
| Refinement (Phenix)<sup>#</sup> |  |  |  |
| Protein residues |  |  | 3776 |
| EMRinger Score |  |  | 2.41 |
| R.m.s. deviations |  |  |  |
| Bond lengths (Å) |  |  | 0.014 |
| Bond angles (°) |  |  | 1.6 |
| Validation |  |  |  |
| Molprobity score |  |  | 1.19 |
| Clash score |  |  | 1.83 |
| Favored rotamers (%) |  |  | 97.08 |
| Ramachandran |  |  |  |
| Favored regions (%) |  |  | 96.32 |
| Disallowed regions (%) |  |  | 0.05 |

<sup>\$</sup> Resolutions are reported according to the FSC 0.143 gold-standard criterion

**Figure S1.**
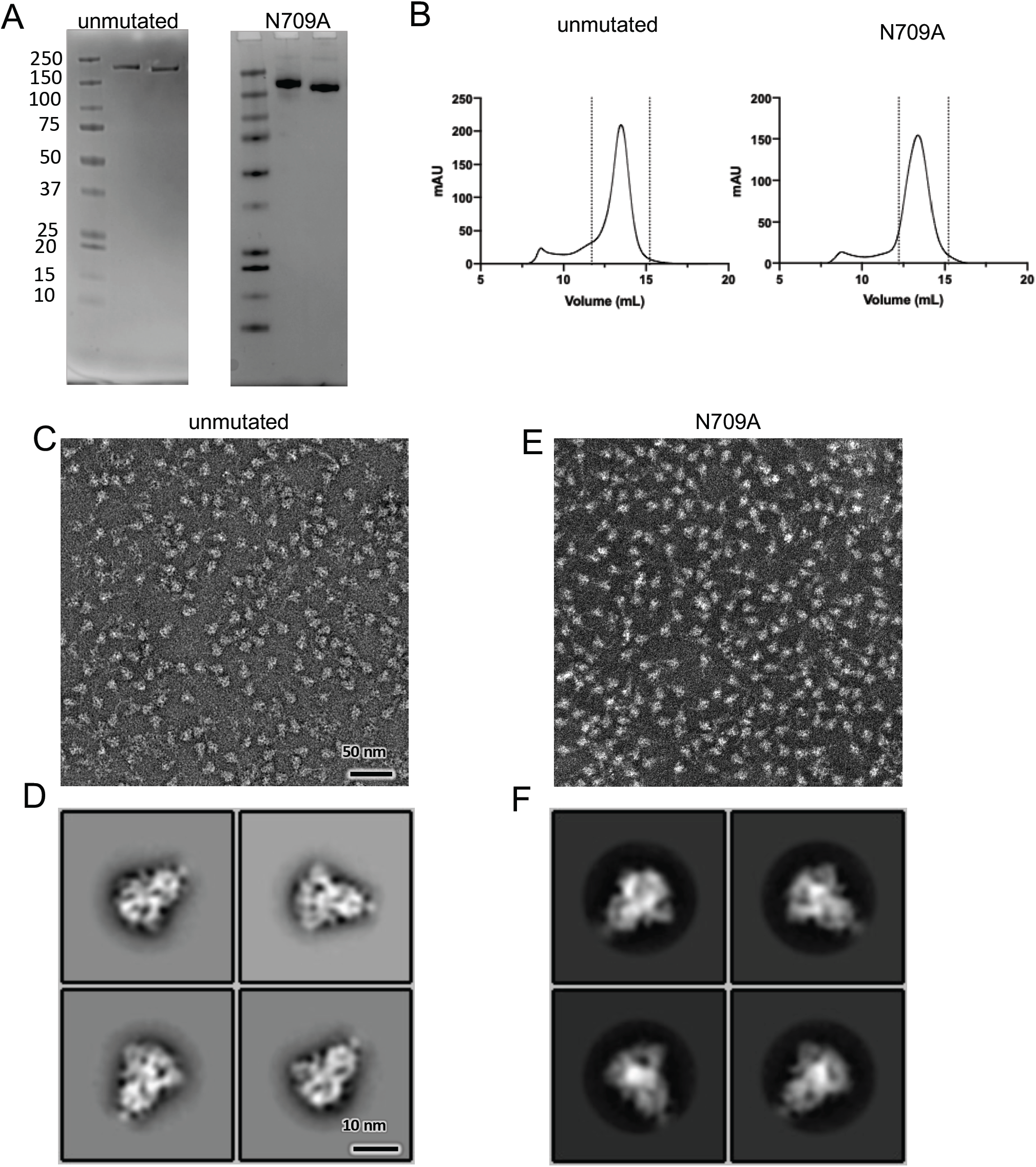
Purification and validation of SARS-CoV-2 S glycoprotein. **(A)** SDS-PAGE analysis of SARS-CoV-2 unmutated (left) and N709A (right) spike. Lane 1: Molecular weight marker; lane 2: elution from StrepTactin resin run under reducing conditions; lane 3: elution from StrepTactin resin run under non-reducing conditions. **(B)** Size-exclusion chromatogram of the affinity-purified SARS-CoV-2V S protein. Data from a Superose 6 10/300 increase column are shown. Dotted lines indicate the fractions that were taken for downstream applications. **(C)** Representative Negative-stain EM (NSEM) micrograph and (D) Representative NSEM 2D class averages of the SARS-CoV-2 unmutated S. **(E)** Representative NSEM micrograph and **(F)** Representative NSEM 2D class averages of the SARS-CoV-2 S with the N709A mutation.

**Figure S2.**
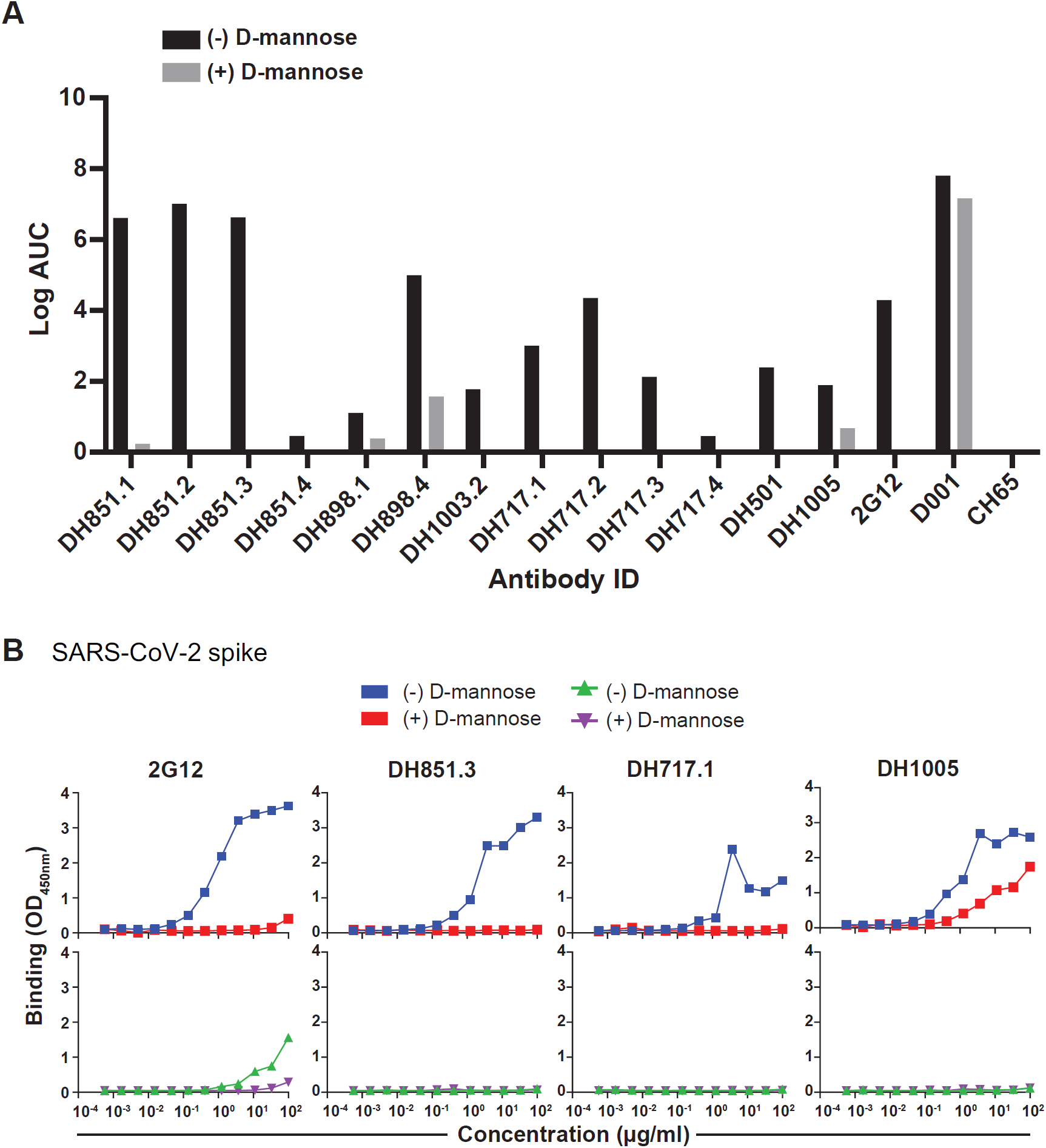
Glycan-dependent binding of FDG antibodies to SARS-CoV-2 spike protein. **(A)** Binding antibody titers reported as Log area under the curve (AUC) for recombinant mAbs tested against SARS-CoV-2 spike protein (See also Figure 1). A fresh aliquot of frozen SARS-CoV-2 spike protein was thawed at 37°C and stored at room temperature overnight prior to ELISA. Spike protein was captured using streptavidin to evaluate FDG mAb binding. Antibody binding was assessed in the absence (-) or presence (+) of D-mannose [1M], and antibody titers were calculated using Softmax or GraphPad Prism software. Shown are binding titers for FDG mAbs, 2G12 (G1M17 version), SARS-CoV-1 RBD (D001) and influenza HA (CH65) mAbs. **(B)** Binding of 2G12 (G1M17 version) and representative FDG mAbs to the SARS-CoV-2 spike ectodomain. Experimental setup was similar to the one described in (A). Blue and red symbols indicate binding in the absence and presence of D-mannose [1M], respectively, from a representative ELISA using BSA-based buffers (see methods). Green and purple symbols indicate binding in the absence and presence of D-mannose [1M], respectively, from a representative ELISA using Superblock or goat serum-based buffers (see methods). Binding was measured at absorbance of OD_450nm_.

**Figure S3.**
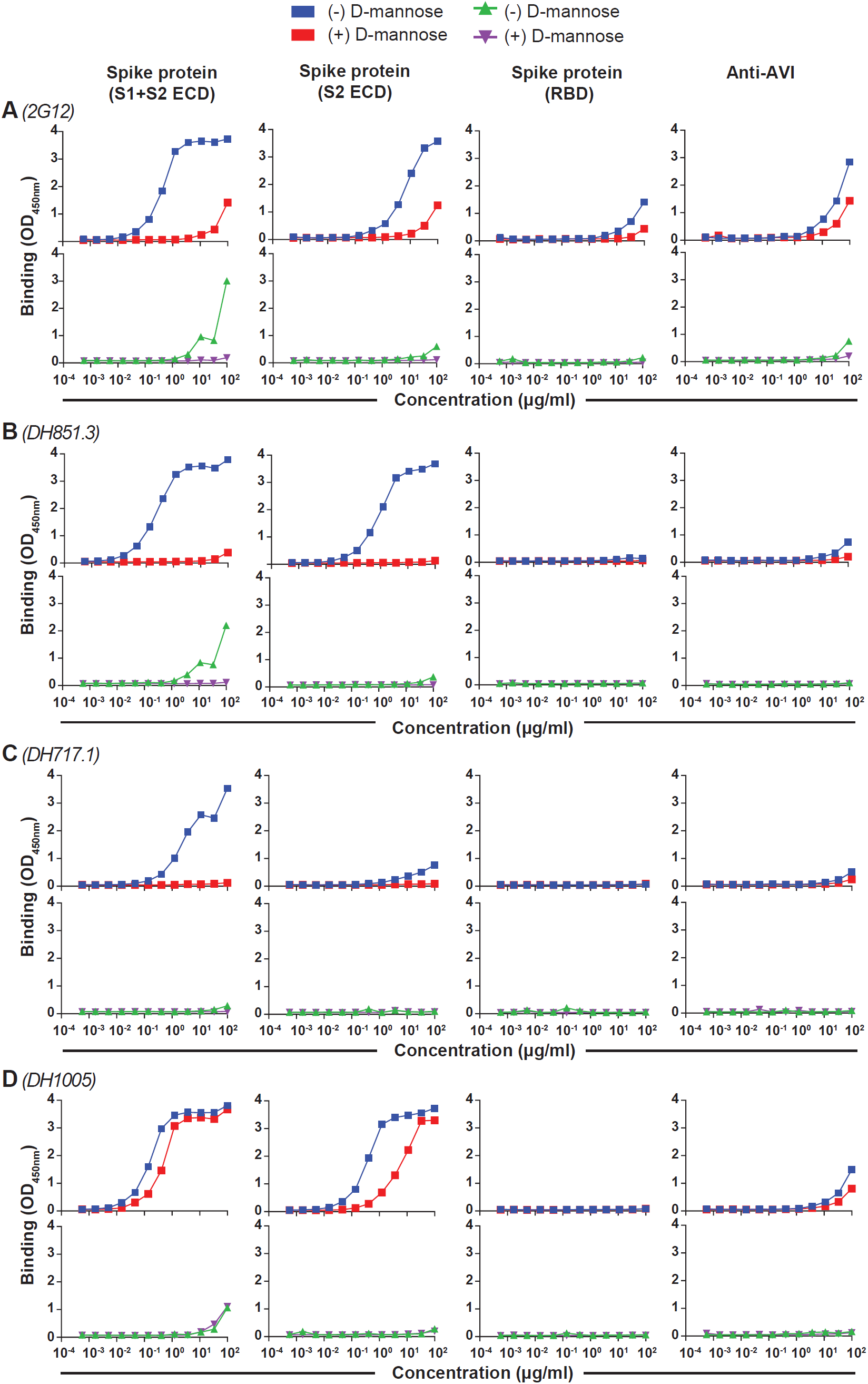
ELISA binding curves of FDG antibodies. **(A-D)** Recombinant FDG mAbs and SARS-CoV-2 spike protein were tested in ELISA. All mAbs were tested for binding in the absence (-) or presence (+) of D-Mannose [1M]. Blue and red symbols indicate binding in the absence and presence of D-mannose [1M], respectively, from a representative ELISA using BSA-based buffers (see methods). Green and purple symbols indicate binding in the absence and presence of D-mannose [1M], respectively, from a representative ELISA using Superblock or goat serum-based buffers (see methods). Commercially available constructs expressing the SARS-CoV-2 S1 and S2 extracellular domain (S1+S2 ECD), S2 domain (S2 ECD), and the receptor binding domain (RBD) were tested for binding by serial dilutions of mAbs; all mAbs were tested at a starting antibody concentration of 100µg/ml. Commercially available proteins were captured using mouse anti-AVI-tag mAb, so we also assessed background binding to only anti-AVI-tag coated plates. Binding levels are shown as absorbance at OD_450nm_ from a representative assay.

**Figure S4.**
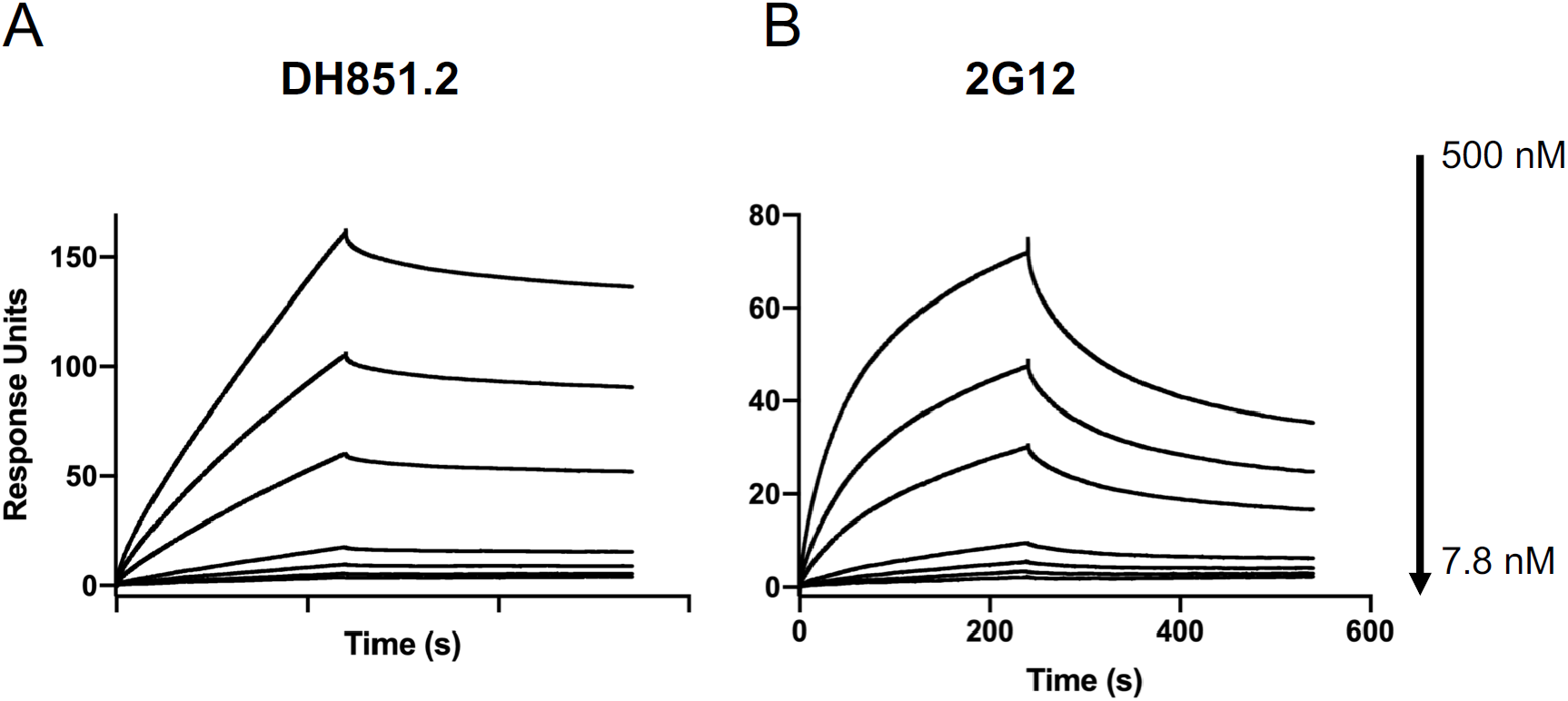
SPR binding curves of DH851.2 IgG and 2G12 IgG. **(A)** DH851.2 IgG and **(B)** 2G12 IgG were tested for binding to the SARS-CoV-2 spike ectodomain in an SPR assay. The SARS-CoV-2 spike at 250nM was captured on a flow cell of an SA chip and binding was measured by flowing over solutions FDG antibodies at 7.8, 15.63, 31.25, 62.5, 125, 250, 500nM in running buffer. The surface was regenerated between injections by flowing over 1M NaCl in 50mM NaOH solution for 10s with flow rate of 100µl/min. Blank sensorgrams were obtained by injection of the same volume of HBS-EP+ buffer in place of IgGs and Fab solutions. Sensorgrams were double-referenced by first subtracting the signal from the reference flow cell and then subtracting the reference-corrected buffer blank.

**Figure S5.**
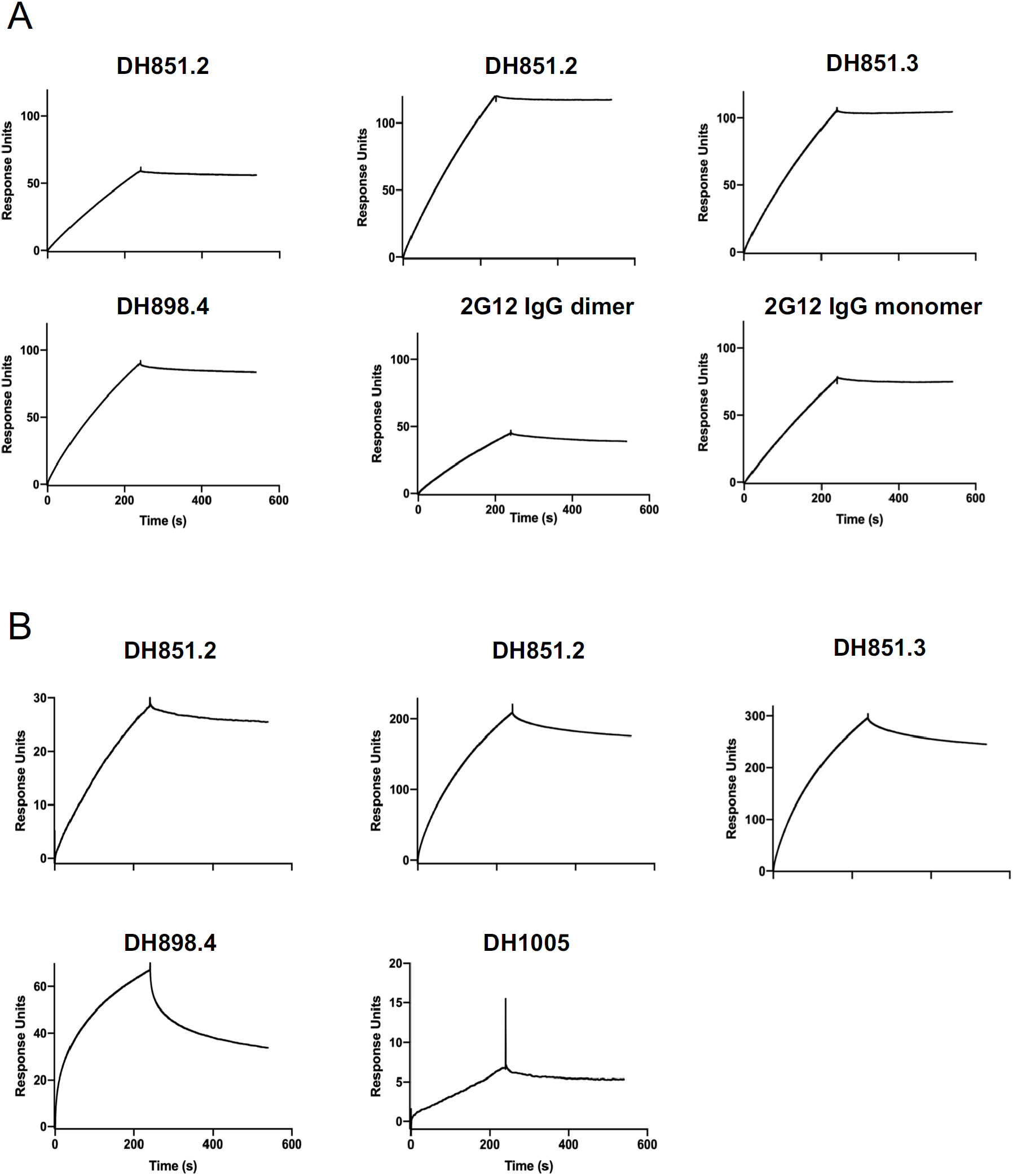
SPR binding curves of FDG antibodies binding to the SARS-CoV-2 spike measured using two different assay formats. The binding of FDG antibodies to SARS-CoV-2 spike was assessed by surface plasmon resonance on Biacore T-200 (GE-Healthcare) at 25°C with HBS-EP+ (10 mM HEPES, pH 7.4, 150 mM NaCl, 3 mM EDTA, and 0.05% surfactant P-20) as the running buffer. **(A)** IgGs were captured on flow cells of a CM5 chip immobilized with human Anti-Fc antibody (8000RU). 200 nM solution of the SARS-CoV-2 spike was flowed over the flow cells. The surface was regenerated between injections by flowing over 3M MgCl2 solution for 10s with flow rate of 100µl/min. **(B)** The SARS-CoV-2 spike was captured on a SA chip, and 200 nM solutions of the IgGs were flown over the flow cells. The surface was regenerated between injections by flowing over 1M NaCl in 50mM NaOH solution for 10s with flow rate of 100µl/min. Blank sensorgrams were obtained by injection of the same volume of HBS-EP+ buffer in place of IgG or SARS-CoV-2 spike solutions. Sensorgrams were double-referenced by first subtracting the signal from the reference flow cell and then subtracting the reference-corrected buffer blank.

**Figure S6.**
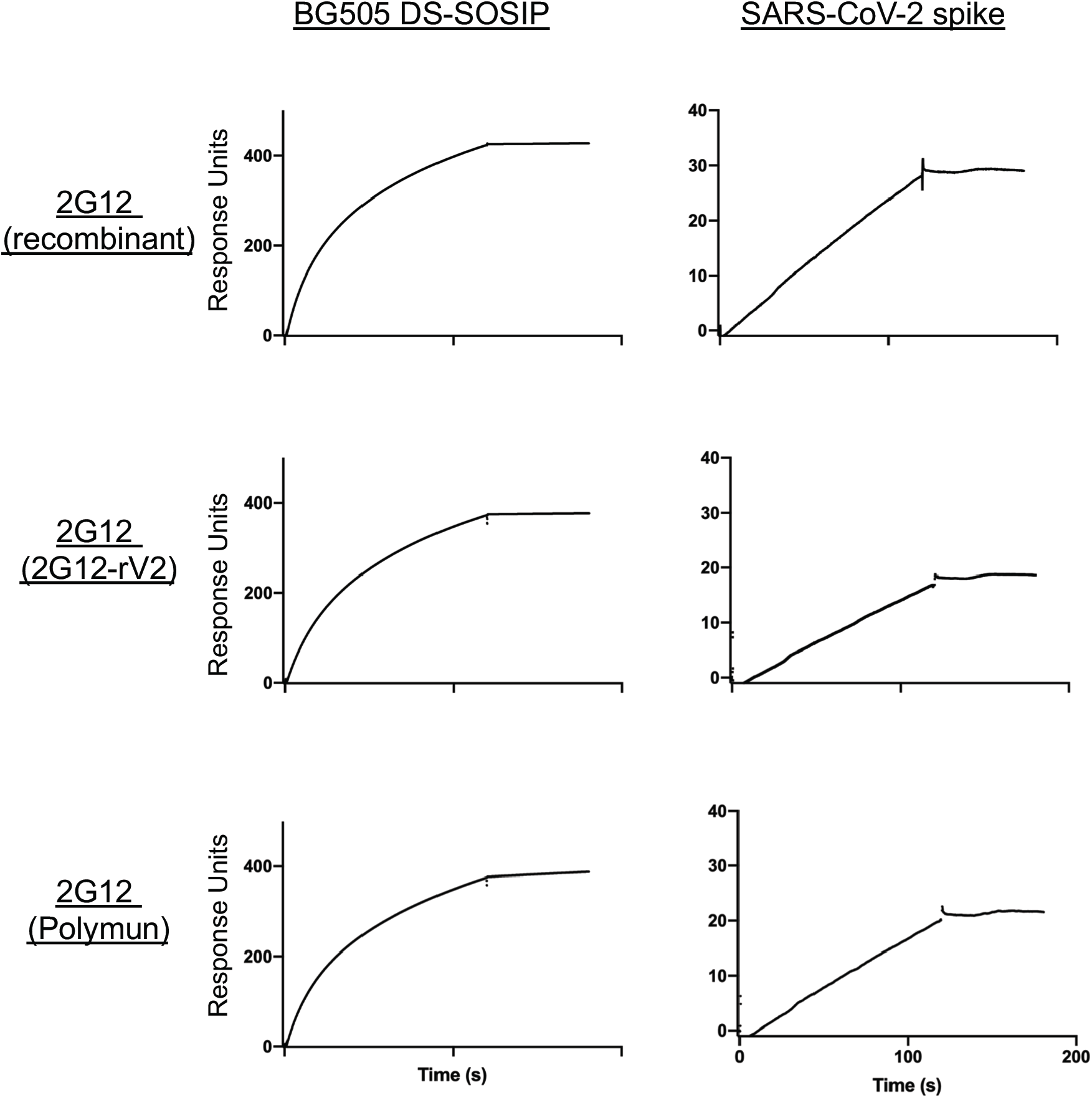
Binding of 2G12 IgG from different sources to HIV-1 Envelope and SARS-CoV-2 spike ectodomain. 200 nM of 2G12 IgG were captured on flow cells coated with anti-Fc antibody. Binding to S protein was assessed by flowing over 200 nM of S ectodomain over the sensor surfaces. The surface was regenerated between injections by flowing over 3M MgCl2 solution for 10s with flow rate of 100µl/min. Blank sensorgrams were obtained by injection of the same volume of HBS-EP+ buffer in place of IgG or SARS-CoV-2 spike solutions. Sensorgrams were double-referenced by first subtracting the signal from the reference flow cell and then subtracting the reference-corrected buffer blank.

**Figure S7.**
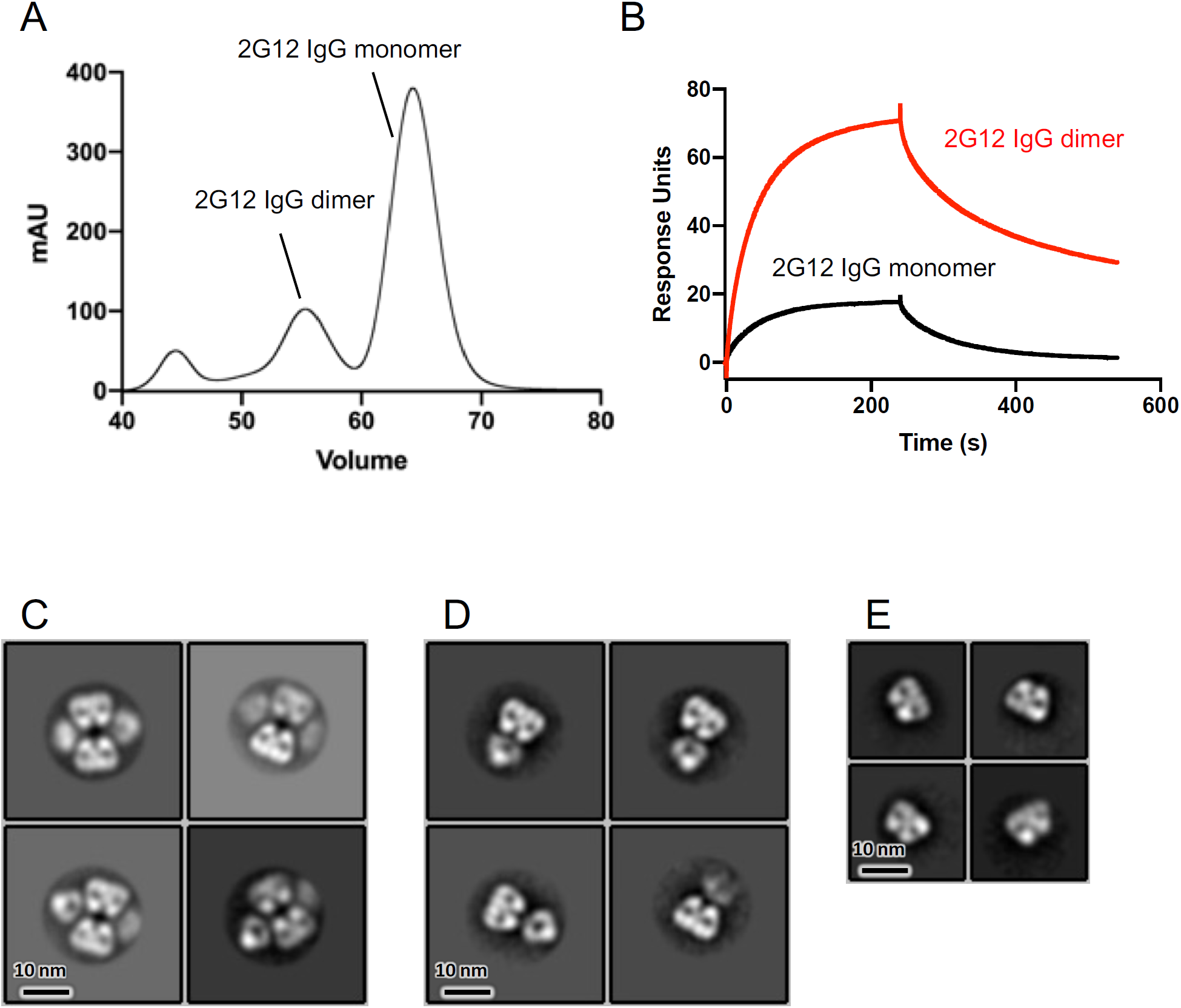
Purification of 2G12 IgG and binding to SARS-CoV-2 S protein. **(A)** Size-exclusion chromatogram of the protein A affinity purified 2G12 IgG. Data from a Superdex 200 16/60 column are shown. **(B)** Surface plasmon resonance sensorgrams showing binding of 2G12 IgG dimer (red line) and 2G12 IgG monomer (black line) to the SARS-CoV-2 S protein. The S protein was captured, via a C-terminal Twin Streptactin tag, on a Streptavidin coated chip and binding was measured by flowing over 1000 nM of the IgG solutions in running buffer. Binding sesnorgrams were blank subtracted and double-referenced. **(C-E)** NSEM 2D class averages of **(C)** 2G12 IgG dimer, **(D)** 2G12 IgG monomer and **(E)** 2G12 Fab obtained by digesting 2G12 IgG monomer with papain.

**Figure S8.**
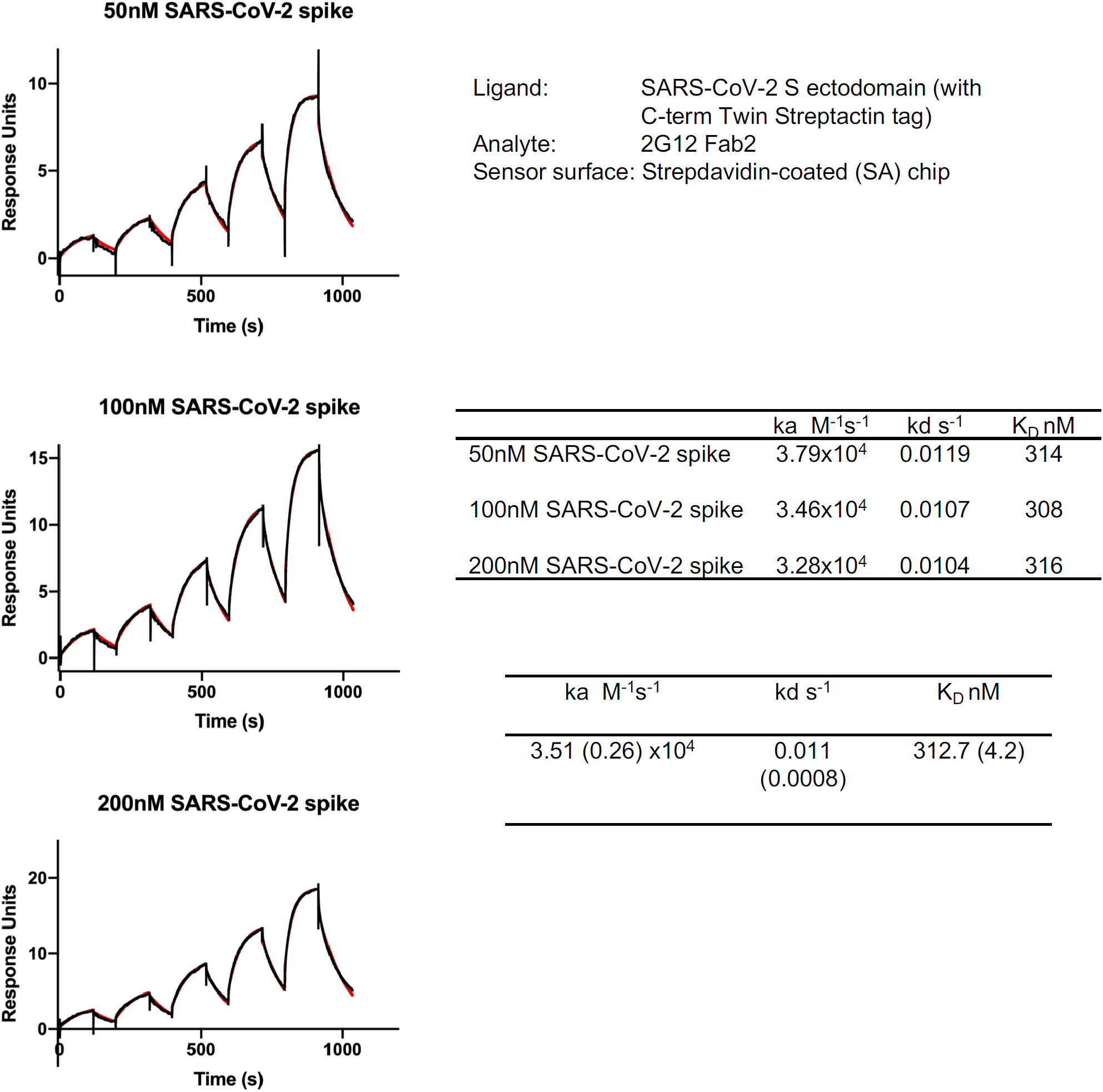
Kinetics and affinity of 2G12 Fab2 binding to SARS-CoV-2 S protein. Graphs show SPR sensorgrams for binding of 2G12 Fab2 to SARS-CoV-2 S ectodomain. Affinity and kinetics of interaction were measured using single cycle kinetics. The experiment was simultaneously run over three surfaces with different levels of spike captured by running over 50 nM (top), 100 nM (middle) and 200 nM (bottom) of SARS-CoV-2 S ectodomain solution. Tables show affinity and kinetics measure for (top) each run separately and (bottom) the mean of the three runs with standard errors shown in parenthesis.

**Figure S9.**
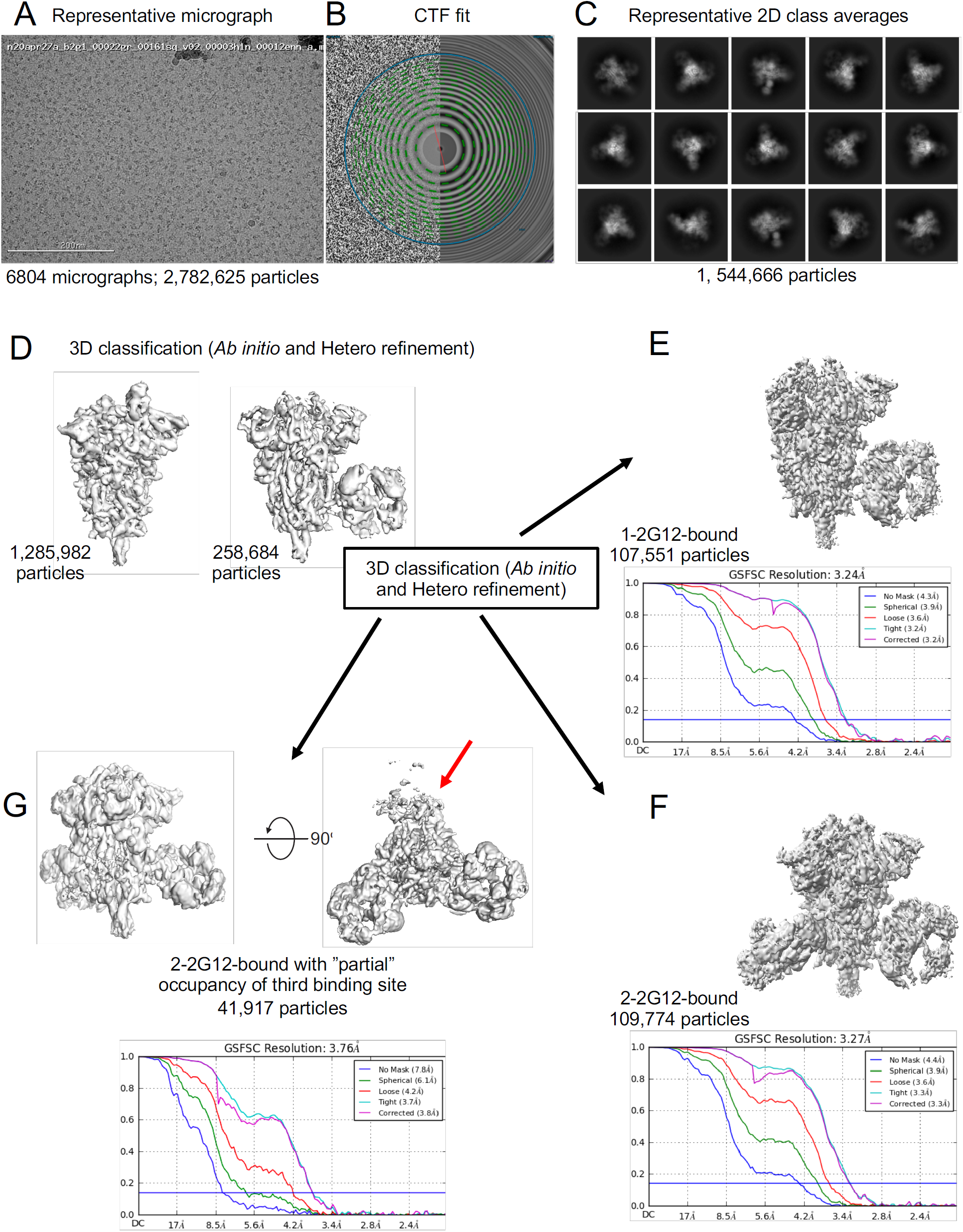
Cryo-EM data processing details. **(A)** Representative micrograph. **(B)** CTF fit **(C)** Representative 2D class averages. **(D)** Maps for unliganded (left) and 2G12-bound (right) S obtained after 3 D classification. **(E-F)** Refined maps for SARS-CoV-2 S protein bound to **(E)** 1 2G12, **(F)** 2 2G12-bound, and **(G)** 3 2G12 Fab2 molecules. Red arrow in **(G)** points to disordered 2G12 Fab2 bound at the third binding site.

**Figure S10.**
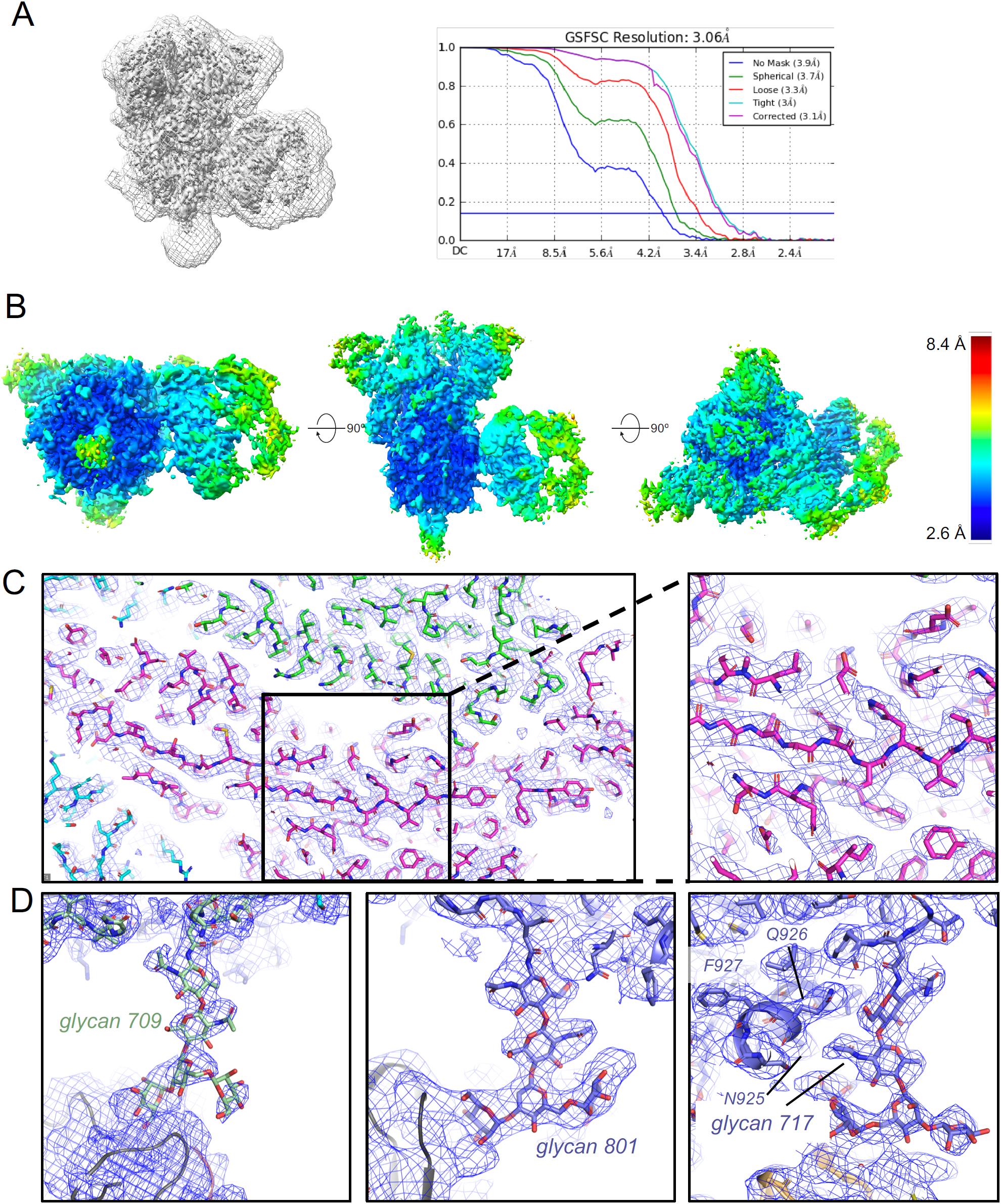
Resolution of cryo-EM reconstruction. **(A)** (Left) Map combining all particles and focusing refinement on the region within the masks that is shown as a grey mesh overlaid on the final refined map shown as a grey surface. (Right) Fourier shell correlation curves. **(B)** Cryo-EM reconstruction of 2G12 bound to the SARS-CoV-2 spike colored by local resolution. **(C)** (left) View of a region in the S2 domain with map shown as blue mesh and fitted model shown as sticks. (right) Zoomed-in view of region shown within the black square in. **(D)** Zoomed-in view of (left) glycan 709, (middle) glycan 801 and (right) glycan 717 bound to 2G12. While not in direct contact with the bound antibody, the HR1 helix may play an indirect role in the binding by stabilizing glycan 717 via a stacking interaction with residues N925 and Q926.

**Figure S11.**
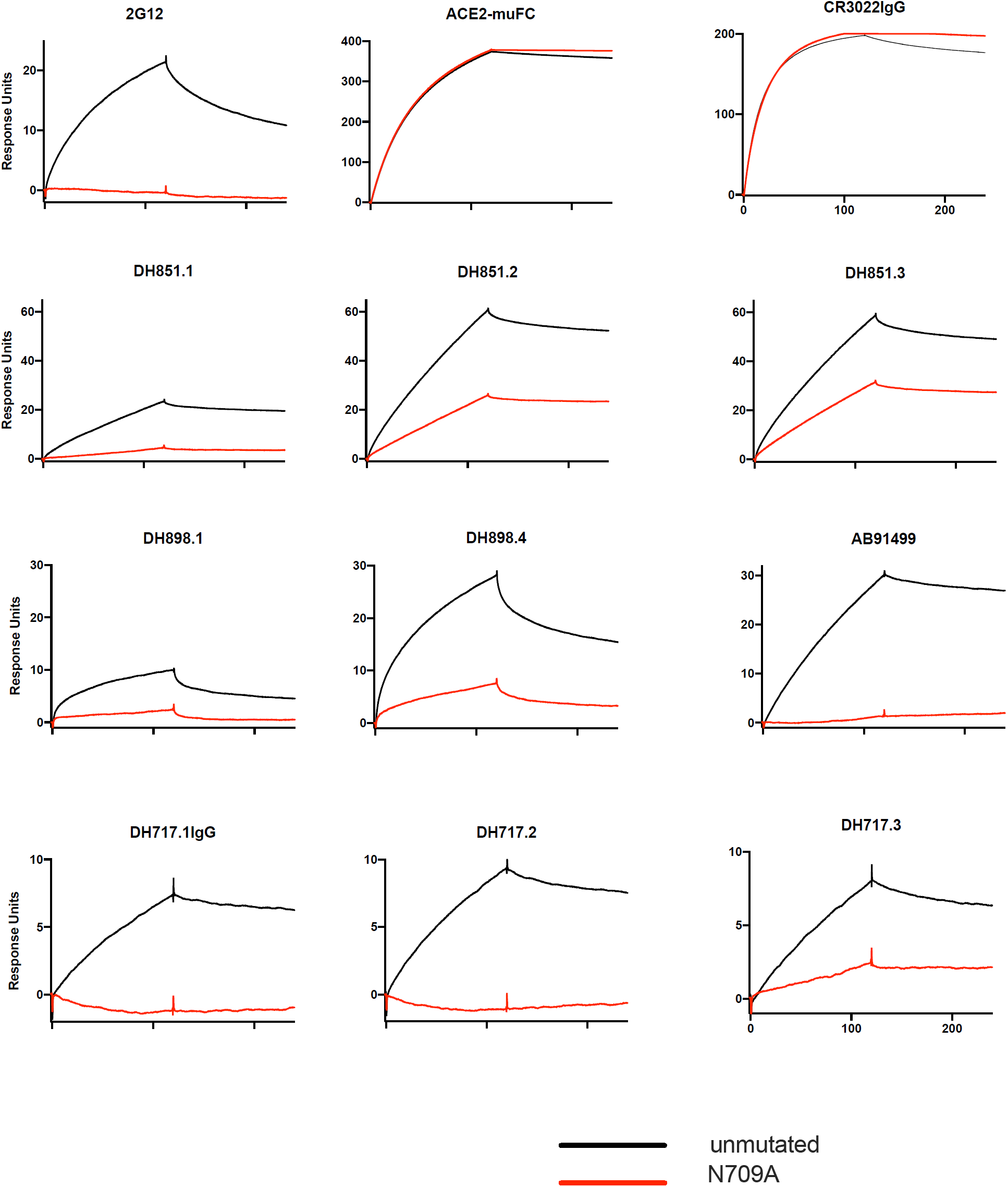
Binding of FDG antibodies, ACE-2 and CR3022 to unmutated and N709-glycan-deleted SARS-CoV-2 spike. Binding between the spikes and a FDG antibody was assessed by SPR by capturing the unmutated spike and the N709-glycan deleted spike on flow cells 2 and 4 of a streptavidin coated (SA) chip, and flowing over a 200 nM solution of each antibody simultaneously over all four flow cells. Flow cells 1 and 3 were used as reference flow cells for flow cells 2 and 4, respectively. Buffer blanks were run in a similar manner and the sensorgrams were double-referenced by first subtracting the signal from the reference flow cell and then subtracting the reference-corrected buffer blank. CR3022 IgG and ACE-2 tagged with a mouse-Fc region were used as controls.

